# *MetaHarmonizer*: robust biomedical metadata harmonization and a contamination control for inflated LLM performance on public benchmarks

**DOI:** 10.64898/2026.06.13.732088

**Authors:** Changchang Li, Abhilash Dhal, Kai Gravel-Pucillo, Kaelyn Long, Michele Waters, Ino de Bruijn, Sean Davis, Sehyun Oh

## Abstract

Public biomedical repositories hold substantial reuse potential, but inconsistent metadata routinely blocks integration across studies. Recent LLM-based harmonization approaches address scale but suffer from non-determinism, hallucinated ontology terms, and, in their highest-accuracy configurations, dependence on proprietary APIs or labeled fine-tuning data. A more fundamental concern is that LLM accuracies on widely-used public benchmarks may substantially inflate transferable capability: under a contamination-controlled evaluation protocol we developed, the apparent LLM-only advantage on the GDC schema-mapping benchmark is inverted and three out of five LLMs recovers 80-100% of GDC identifiers from zero-schema context, suggesting direct memorization. Building on this insight, we present *MetaHarmonizer*, an automated metadata harmonization system designed to be robust by construction: *SchemaMapper* aligns attribute names across schemas, and *OntologyMapper* standardizes values to controlled vocabularies. Both modules implement a multi-stage cascade that escalates to more resource-intensive methods only when earlier stages fall short, with all candidates grounded in pre-defined controlled vocabularies to preclude hallucinated outputs and LLMs used only as bounded preprocessing components rather than inference-time dependencies. On the GDC schema-matching benchmark, *SchemaMapper* with the deployment-optimized LLM-generated alias dictionary achieved 71.6% Top-1 accuracy and the higher Recall@GT than Magneto bipartite variants, recovering significantly more ground-truth mappings; with the best performing alias dictionary, it reached the highest Top-1/Top-5/Recall@GT, and also matched the best Magneto reranker (fine-tuned LLM-reranker) on MRR; and it also outperforms LLM-only performance under contamination-controlled conditions. On four EFO benchmarks, *OntologyMapper* achieved 77.9–95.5% Top-1 accuracy, outperforming *text2term* by up to 16.4 pp and direct LLM inference (against the smaller corpus) by 19.2 pp because memorization is not a viable shortcut for this task. Across both modules, calibrated confidence scores separate correct from incorrect predictions (AUC 0.73–0.94), enabling principled human-in-the-loop triage. Inference is fully local, deterministic, and computationally efficient – seconds on schema mapping and under a minute for ontology mapping of up to ∼7,000 terms against the pre-indexed 33,230-term corpus. Released as a Python package with a domain-agnostic architecture, *MetaHarmonizer* provides a scalable foundation for improving the FAIRness of biomedical data and enabling cross-study integration, alongside an evaluation methodology applicable to any LLM-augmented bioinformatics benchmark built on public benchmarks.

## Introduction

The exponential growth of multi-omics data has transformed our understanding of biological systems and disease mechanisms. However, the utility of these vast datasets depends heavily on the quality and accessibility of their associated metadata, which provides crucial context on experimental conditions, patient characteristics, and treatment outcomes. While significant attention has been paid to standardizing and sharing omics data^1–3^, harmonizing patient- and sample-level metadata remains a critical yet often overlooked challenge in the biomedical research ecosystem.

Metadata from public biomedical repositories often exhibits inconsistent terminology, structure, and completeness. This heterogeneity severely impacts the FAIRness (Findability, Accessibility, Interoperability, and Reusability) of valuable research data, making it challenging to perform cross-study analyses or integrate datasets from multiple sources. The situation is particularly difficult for patient/sample-level metadata, where variations in naming conventions, units, and formatting create significant barriers to data identification, integration, and reuse^4,5^.

Computational approaches to metadata harmonization can be organized along two levels: *schema-level alignment*, which maps variable names or column definitions across datasets, and *value-level ontology grounding*, which maps free-text values within those variables to controlled vocabulary terms. The two tasks pose different challenges and have been addressed by largely separate lines of work.

Methods for matching variable names or column definitions have progressed from lexical similarity to neural and embedding-based approaches. Mallya et al.^6^ reported 98.95% Top-5 accuracy using fully convolutional networks with BERT embeddings for cardiovascular variables, although the task was framed as paired-sentence classification across three datasets with rich semantic descriptions rather than open-vocabulary alignment. Salimi et al.^7^ showed that LLM embeddings could support schema harmonization across Parkinson’s disease studies, reaching average accuracies above 80%. At the level of phenotype data elements, Pathak et al.^8^ found that 40% still required manual curation, underscoring the residual burden even when computational support is available. Most relevant to the present work, Liu et al.^9^ introduced Magneto, which combines small language models for candidate retrieval with LLM-based reranking and was evaluated on a benchmark of 736 GDC data dictionary attributes; its best-performing variant requires LLM API calls at inference time and depends on domain-specific fine-tuning data.

A larger body of work has tackled the more heterogeneous task of mapping free-text values to ontology terms, where lexical and rule-based methods remain the most widely used strategy. Miotto et al.^10^ processed over 90,000 influenza records using pattern matching, demonstrating early feasibility for structured extraction. Urbanowicz et al.^11^ reported ∼85.5% mapping rate through automated exact string matching, and Mate et al.^12^ processed 98.7% of records across 10 European Biobanks. Gonçalves et al.^13^ developed *text2term*, an open-source tool that maps free-text biomedical entity descriptions to ontology terms using Term Frequency-Inverse Document Frequency (TF-IDF)-based cosine similarity and edit distance, achieving 73–81% Top-1 accuracy on curated EFO benchmarks. Hybrid pipelines that incorporate ontology resources extend coverage further; Grossman et al.^14^ reached 94.3–99.6% using MetaMap^15^ and UMLS, but require substantial domain expertise for rule curation and incur an ongoing maintenance burden as terminologies evolve. These studies consistently report failures with typographic variation, ambiguous abbreviations, and non-standardized naming, and plateau at 60–80% automation, leaving a substantial manual harmonization burden.

Embedding- and LLM-based approaches have begun to close this gap recently. Dylag et al.^16^ used Sentence-BERT^17^ clustering to achieve a 117-fold speed improvement over manual curation, but at the cost of moderate accuracy (V-measure 0.237) and limited validation. Higashi et al.^18^ combined LLMs with semantic clustering to harmonize four key attributes across more than 400,000 human gut microbiome samples, and Ikeda et al.^19^ applied LLM-based methods to extract biological terms from over 40 million epigenomics records. The strengths and limits of LLM-based grounding are now well documented: Verbitsky et al.^20^ standardized 691,220 terms across three clinical domains and reported 90–96% accuracy for in-dictionary terms but only 12–17% for out-of-dictionary terms.

Looking across these efforts, several systemic limitations remain unresolved. First, most studies remain confined to a single domain, data type, or repository, and cross-domain evaluation is largely absent. Second, LLM-based approaches, while scalable, still require substantial human oversight and face fundamental challenges: non-deterministic outputs that conflict with FAIR principles^21^, hallucination of plausible but nonexistent ontology terms^22^, limited contextual awareness across thousands of variables^23^, and practical barriers including API costs and data privacy concerns^24^. Third, most approaches address only one facet of the harmonization problem, leaving the full spectrum of curation tasks required for cross-study integration unaddressed, including reconciling conflicting values, consolidating redundant metadata fields, and incorporating study-level context. The accuracy–interpretability tradeoff remains unresolved: high-accuracy neural approaches operate as black boxes, while interpretable lexical methods plateau at insufficient levels of automation. No existing system adequately balances automation completeness, cross-domain robustness, computational efficiency, and interpretability.

To address these limitations, we developed *MetaHarmonizer*, an automated metadata harmonization framework comprising two modules: *SchemaMapper* for column-level alignment and *OntologyMapper* for value-level ontology grounding. *MetaHarmonizer* implements a progressive, multi-stage architecture that balances speed, cost, and accuracy. The framework combines hybrid embedding strategies grounded in real ontology terms with calibrated confidence scores that enable practical human-in-the-loop workflows. Benchmarked against the most directly comparable tools (*Magneto* for schema matching and *text2term* for ontology mapping), *SchemaMapper* achieves competitive or better performance with substantially lower computational cost and no dependence on external LLM APIs at inference time, and *OntologyMapper* displayed superior performance on all metrics used. We further show that *OntologyMapper* outperforms LLM-only inference, and that, under contamination-controlled evaluation, *SchemaMapper* paired with an LLM-generated alias dictionary matches LLM-only performance without incurring inference-time API costs. In the course of this benchmarking, we developed a contamination-control protocol (combining source-side paraphrase, target-side identifier renaming, and direct memorization probes) that quantifies the extent to which LLM-only performance on public schema benchmarks reflects pretraining exposure rather than transferable matching capability. We present this methodology alongside *MetaHarmonizer* as a general-purpose contamination control for LLM-augmented benchmarks.

## Methods

### Architecture

*MetaHarmonizer* comprises two modules that address complementary harmonization tasks: *SchemaMapper* aligns heterogeneous attribute names to a target schema, and *OntologyMapper* standardizes free-text values to controlled vocabulary terms. Both modules implement a multi-stage cascade that escalates from inexpensive exact and lexical matching to embedding-based methods, with all candidates drawn from pre-defined controlled vocabularies; the sections below describe each stage in detail. Within this design, *stages* are sequential steps in the pipeline, and *methods* are alternative or sub-approaches available within a stage. *MetaHarmonizer* is implemented in Python, with FAISS-based vector search, SQLite-based caching, and connectors to external terminology services (NCI, UMLS, OLS).

### Prepare Inputs

There are two required inputs for *MetaHarmonizer*: 1) Queries: column names for *SchemaMapper* and values for *OntologyMapper* that are subject to harmonization, and 2) Targets: a list of target attributes for *SchemaMapper* and a corpus of allowed ontology labels for *OntologyMapper*. Both column names and values are normalized through case normalization, whitespace handling, and removal of special characters. During the input pre-processing for *SchemaMapper*, non-informative values (i.e., *yes/no/unknown*) are also removed from the list of unique values; while they have information on the attribute’s status (denoted by the column name), they do not provide any additional information during Stage 2 of the schema mapping pipeline. Removing these values results in the special case (*M* = 0) during Stage 2-2 of *SchemaMapper*. The key parameters for each module are listed in **Supplementary Tables 1-2**.

### SchemaMapper

*SchemaMapper* has a cascade architecture with 1) early termination when confidence is high, avoiding unnecessary computation, 2) best-so-far tracking, ensuring a result is always returned even if no threshold is met, and 3) fine-grained control through per-strategy thresholds. We set the default thresholds based on empirical tests using manually created gold-standard datasets^25^ distinct from the benchmark datasets.

#### Stage 1. Exact/Fuzzy

This stage performs exact and fuzzy matching between the queried column names and the target schema’s attribute names. For fuzzy matching, we used the token_sort_ratio function from the *RapidFuzz* library^26^. There are four method tags for exact (exact) vs. fuzzy (fuzzy) matching against target (std) vs. its alias (alias); std_exact, alias_exact, std_fuzzy, and alias_fuzzy. Aliases can be sourced from manual curation or LLM-based generation (see the section, ‘LLM-generated alias dictionary for *SchemaMapper*’, below).

#### Stage 2. Column Value Level

This stage is invoked when the query includes column values. It targets unmatched, non-numeric columns from Stage 1 and uses their unique, non-null data values as inputs. Each value is embedded individually and compared against the embeddings of allowed values from a ‘target attribute + allowed values’ dictionary. For each query, Top-k similar dictionary values vote for their parent field; votes are aggregated per target field using log-compressed frequency weights, and a field is returned when its weighted proportion exceeds the threshold (default: 0.2).

##### 2-1. Allowed Value Matching (method = ‘value’)

This method applies to columns containing a *relatively stable set of categorical values*, such as sex, country, and ethnicity. The algorithm defines a ‘hit’ as any value pair above the threshold (default is cosine similarity >= 0.85). The ‘hit rate’ is calculated for each input column *i* as:

*d_i_* = number of unique values in input column *i*
*n_i_* = number of input values that achieve ‘hit’ status (cosine similarity >= 0.85 with any target value)
*p_i_* = hit rate

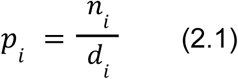

In the special case where *d_i_ = 0* (e.g., values are limited to *yes/no/unknown* and are removed during preprocessing), we set *p_i_ = 0* and note that there is insufficient evidence. Input columns are ranked by *p_i_*in descending order. When multiple columns have identical *p_i_*, they are ordered by their average similarity scores.

##### 2-2. NCIt Voting Method (method = ‘ontology’)

This method is triggered for columns where possible values are actively changing or growing, such as treatment names and diseases. We map unique values from each column to ontology terms using the NCIt client, leveraging NCIt’s built-in fuzzy matching and synonym/alias expansion. Then we traverse the NCIt hierarchy upward to determine whether mapped terms belong to predefined ontology nodes per categories. The examples of predefined categories include ‘Body Part’ (NCIT:C32221) for body_site, ‘Disease or Disorder’ (NCIT:C2991) for disease, and ‘Pharmacologic Substance’ (NCIT:C1909) for treatment_name.

*M* = total number of input values successfully assigned to NCIt terms
*m_i_* = number of values assigned NCIt terms that match the target category *c_i_*
*v_i_* = NCIt voting score/proportion for category *c_i_*

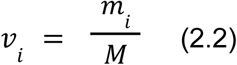

If *M* = 0 (no values successfully mapped to NCIt terms), then *v_i_* = 0 and the result is noted as ‘no NCIt evidence’. The score *v_i_* represents the proportion of successfully mapped values that belong to the target category *c_i_*, providing a confidence measure for categorizing the input column.

### Stage 3. Column Name Level

This stage contains two methods for different value types (numeric vs. categorical).

#### 3-1. Numeric Column Matching (method = ‘numeric’)

This method is used if a column predominantly contains numeric values, verified by numeric-type checks and unit inference. For example, columns containing numbers followed by “*years/months/days*” are classified as “time” type, while those followed by “*mg/ml/AUC*” are classified as “dose” type. The similarity score for numeric fields combines two components: the primary component is the cosine similarity between sentence transformer embeddings of the query field (*e(q)*) and candidate field *i* (*e(ci)*), and the second component is family boost, an additional factor applied when both query and candidate columns belong to the same type. β_family_ is a weighting parameter that controls the strength of the family boost, and *I* is an indicator function that returns 1 when the query and candidate belong to the same family, and 0 otherwise.

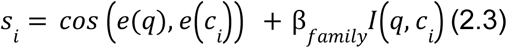

#### 3-2. Semantic Column Matching (method = ‘semantic’)

This stage processes unseen or ambiguous non-numeric column names that failed to match in Stage 2, as well as low-confidence matches from Stage 3-1. The alias dictionary (the same one used in Stage 1) for target columns can be used in this stage as well. It computes semantic similarity between the query and the target/alias column names.

##### LLM-generated alias dictionary for *SchemaMapper*

To improve schema mapping performance, we expand each target field into plausible surface forms provided through an alias dictionary. *SchemaMapper* uses this dictionary both as a string index (for rule-based matching at Stage 1) and as an embedding index (for semantic retrieval at Stage 3). Aliases for each target attribute were generated from five structured passes, each targeting a distinct class of surface variation hinted from the previous observation^25^:

1. Synonym pass: alternative natural-language phrasings (e.g., `vital_status → survival status`).
2. Abbreviation pass: acronyms and abbreviated forms commonly used in clinical and research databases (e.g., `lymph_nodes_examined_positive → LN+, LNs_pos`).
3. Value-encoded boolean indicators (e.g., `ADJUVANT_CHEMO → treatment_type`).
4. Composite pass: fields with embedded modifiers or qualifiers (e.g., PRIOR_, BASELINE_, AGE_AT_).
5. Institutional pass: plausible institution- or consortium-specific variants (e.g., TCGA_SUBTYPE, CPTAC_TUMOR_GRADE).

Each pass operated on batches of 20 fields and returned a structured CSV with columns source, field_name, and is_numeric_field. We tested four Anthropic models via API (`claude-opus-4-7, claude-sonnet-4-5, claude-haiku-4-5, claude-opus-4-5`), two Google API models (`gemini-2.5-flash, gemini-2.5-pro`), and three open-weight models via local Ollama inference (gemma3:27b, gemma4:26b, and qwen3:32b) (**Supplementary Table 9**). Outputs from all five passes were concatenated, deduplicated on the ‘source:field_name’ pair, and screened for hallucinated field names (i.e., aliases generated for fields not present in the input schema) that were removed before downstream use. The alias dictionary is target-schema-specific and requires no labeled training data; only the target field names are needed to generate it.

The alias dictionary augments two schema-matching stages: In Stage 1, *SchemaMapper* performs a normalized lookup over the alias index and applies *RapidFuzz* token-sort-ratio matching against the alias set when no exact hit is found. This is cascaded immediately after its standard dictionary counterparts. In Stage 3, *SchemaMapper* fuses Top-*k* retrievals from the standard field-name index with Top-*k* retrievals from the alias index, using pre-computed sentence-transformer embeddings of all alias strings. Stage 2 (value-based) does not consume the alias dictionary.

##### OntologyMapper

Our ontology mapping module employs a progressive multi-stage pipeline, where each stage handles increasingly complex matching scenarios. Once a term is resolved at a given stage, it is removed from the candidate pool and not re-evaluated by downstream stages.

#### Index Construction

We prebuild a local ontology term corpus from selected root ontology terms. The corpus contains all descendants of the root, including label, OBO ID, description, and synonyms. From this corpus, we construct per-category SQLite tables used by Stage 2.5; terms are fetched once per category from the NCI EVS REST API (for NCIt) or EBI OLS4 API (for non-NCIt). Alternatively, the tables can be populated offline from a pre-saved corpus JSON. A paired FAISS index is then built over the synonym rows, so each FAISS vector ID equals its SQLite row ID.

##### Stage 1. Exact Matching

This stage attempts a direct lookup of each query against the ontology corpus without expansion.

##### Stage 2. Semantic Matching

Queries unresolved by Stage 1 are passed to Stage 2, which performs vector retrieval over the same ontology corpus. Two preprocessing steps are applied to the residual queries before embedding, both implemented as query-side transformations - lightweight normalization and shortname-to-fullname expansion. The transformed queries and the corpus are embedded with a biomedical encoder. We use SapBERT^27^ as the default model in this stage. SapBERT is primarily trained on the Unified Medical Language System (UMLS), a massive metathesaurus and collection of biomedical ontologies containing over 4 million concepts and 10 million+ synonym pairs. We implement two alternative methods (the default is LM) for generating text representations. *OntologyMapper* provides two interchangeable Stage 2 backends: a sentence-transformer pipeline (ST) that uses mean-pooled token embeddings, and a language-model pipeline (LM) that uses CLS-token embeddings (no statistically significant difference was observed in the tests conducted in this study). In both cases, query and corpus vectors are L2-normalized, and the Top-k corpus entries are retrieved per query by cosine similarity. Each candidate is returned with its similarity score in [0, 1] and assigned a match level of 2.

##### Stage 2.5. Synonym Dictionary Boost

Stage 2.5 targets results from Stage 2 whose Top-1 scores fall below the threshold (default: 0.9). For each such query, it retrieves Top-k matches from an FAISS-indexed ontology synonym dictionary, merges them with the Stage 2 candidates (taking max score per duplicate), and re-ranks. The Stage 2 row is replaced with the merged ranking only if the new Top-1 score is higher or the curated term reaches a better match level; otherwise, Stage 2 is left untouched.

### SchemaMapper Benchmarking

We benchmarked *SchemaMapper* against Magneto (version: 0.3.0.dev0)^9,28^, an end-to-end schema-matching system. We used four reported Magneto variants: zero-shot (zs) and fine-tuned (ft) versions with bi-encoder (bp) and fine-tuned LLM (llm) re-rankers (Magneto-zs-bp, Magneto-ft-bp, Magneto-zs-llm, and Magneto-ft-llm). We reproduced the Magneto bipartite (BP) graph reranker in both zero-shot (zs) and fine-tuned (ft) configurations using the published code and benchmark data. For the Magneto LLM reranker variants, we report the values from the original paper, as these require access to the proprietary LLM API. The target benchmark data were 10 Clinical Proteomic Tumor Analysis Consortium (CPTAC) studies previously used by Magneto. Each study contains heterogeneous metadata column names, which were manually curated by domain experts into 736 standardized target attributes in the GDC data dictionary. Across the 10 studies, 165 source–target pairs had curated ground-truth annotations^28^. Ground-truth annotations include one-to-many mappings, where a single source column maps to multiple acceptable targets.

### OntologyMapper Benchmarking

We compared *OntologyMapper*’s performance against *text2term* (v4.1.2)^13^, a widely used tool for mapping free-text descriptions of biomedical entities to ontology terms. We used benchmark datasets previously established for *text2term*: (1) The UKBB-EFO *benchmark* (n = 888) consists of mappings between UK Biobank phenotype descriptions and EFO terms, derived from the curated mapping set described by Sollis et al.^29^. This dataset represents the most challenging scenario, as UK Biobank phenotype descriptions are often colloquial, abbreviated, or otherwise divergent from formal ontology labels. (2) The Biomappings-EFO *benchmark* (n = 795) draws from the Biomappings community-curated collection of cross-ontology mappings^30^, filtered to those targeting EFO. These mappings use source ontology term labels as input strings, providing a scenario in which source terms are structured ontology labels rather than free text. (3) The OLS-EFO benchmarks use cross-ontology mappings hosted in the EMBL-EBI Ontology Lookup Service (OLS). We evaluated two subsets: *a disease-restricted subset* (n = 5,770 for *OntologyMapper*; n = 5,824 for *text2term*) and the *full cross-ontology mapping corpus* (n = 7,377 for *OntologyMapper*; n = 7,504 for *text2term*). As with Biomappings-EFO, source terms are ontology labels. Query-count differences between tools reflect divergent handling of rows that share a query string but map to different targets; see ‘Denominator conventions across analyses’ section in the Supplementary Methods for the per-analysis denominator scheme.

For the head-to-head benchmark evaluation, we constructed a merged reference corpus for *OntologyMapper* rooted in the Experimental Factor Ontology (EFO, v3.62.0). We extracted 51,862 native Compact Uniform Resource Identifiers (CURIEs) from the EFO OWL distribution. Restricting to the twelve ontologies that EFO cross-imports (EFO, MONDO, Orphanet, CHEBI, HP, UBERON, CL, GO, OBI, PATO, DOID, and BFO) and flattening imported terms to the EFO namespace yielded 39,460 rows compatible with *OntologyMapper*’s single-source partitioner. The module’s default obsolete filter then removes rows whose label begins with obsolete_, leaving 33,230 live terms used for retrieval. Corpus composition is EFO-dominated (16,218 terms; 48.8%) with substantial contributions from MONDO (9,508; 28.6%), Orphanet (2,051; 6.2%), CHEBI (1,722; 5.2%), and HP (1,591; 4.8%); UBERON, CL, GO, OBI, PATO, DOID, and BFO together account for the remaining 6.4%. Synonyms were extracted in parallel, producing a synonym index of 142,645 rows covering exact, related, narrow, and broad synonym axiom types. We used the same 12-ontology EFO corpus (v.3.62.0) as a target corpus for *text2term*. Due to different parsing strategies, this same target corpus resulted in 33,659 terms (∼1.3% margin); the additional 429 terms are mostly label-less or duplicate entries irrelevant to actual mapping.

### Runtime Analysis

For *SchemaMapper* runtime analysis, each benchmark was timed under two conditions: 1) End-to-end per study wraps a fresh mapping module instantiation – SentenceTransformer load, target-schema encoding, and Stage 1/2/3 retrieval over every source column. 2) Per-column, per-stage records wall-clock time for each (column, stage) call. *SchemaMapper* does not persist an embedding index between runs: target-schema and alias embeddings are recomputed within each module instantiation, so the initialization cost is paid once per study rather than amortized across a disk cache. All timings were collected on a single workstation: Apple M4 Pro, 48 GB unified memory, internal SSD, macOS Sequoia 15.6.1 (aarch64); Python 3.13.5, PyTorch 2.8.0 (CPU-only; the MPS backend was not used), sentence-transformers 5.1.1. Embedding inference was executed on a CPU with single-threaded BLAS (Basic Linear Algebra Subprograms).

Each *OntologyMapper* benchmark was timed under three conditions: 1) *Cold* measures a first-time *OntologyMapper* run at a previously unseen corpus (available as a local OWL file); it includes corpus FAISS embedding, synonym FAISS embedding, and Stage 1/2/2.5 retrieval. 2) *Warm* measures a repeat *OntologyMapper* run that reuses cached FAISS indices and executes Stage 1/2/2.5 retrieval. 3) *text2term* measures a single invocation per benchmark, with ontology parse amortized across benchmarks within a session. Warm runs reuse a three-layer cache: SQLite concept tables, an on-disk FAISS index, and in-process model weights that are populated once per (model, corpus-content) pair. Timings used the same workstation as *SchemaMapper*, with FAISS 1.11.0 and *text2term* v4.1.2 added.

### Performance Evaluation

We adopted four evaluation components.

1. ***Top-k accuracy*** (k = 1,3,5) measures the proportion of queries for which the correct target column appears among the Top-k returned candidates. For queries with multiple acceptable ground-truth targets, we applied the “any-match” criterion, counting a prediction as Top-1 correct if the Top-1 prediction matched any acceptable target.
2. ***Mean Reciprocal Rank (MRR)*** reports the average inverse rank of the first correct match, rewarding systems that place correct matches near the top.
3. ***Recall at Ground Truth size (Recall@GT)***, a standard metric from the Valentine schema matching benchmar^9,31k^ and adopted by Magneto^9,31^, measures the proportion of acceptable ground-truth targets recovered within the Top-G candidates, where G is the number of the ground-truth matches per query; when G = 1, Recall@GT reduces to Top-1 accuracy, and when G > 1, it captures the system’s ability to retrieve all valid correspondences. Unlike Top-k and MRR, which depend only on within-query rankings, Recall@GT admits two non-interchangeable formulations – per-query and global – that differ in whether scores must be calibrated across queries; full definitions and the conditions under which each is computable are given in Supplementary Methods.
4. ***Confidence calibration*** assesses whether confidence scores discriminate correct from incorrect Top-1 predictions, since downstream human-in-the-loop review depends on calibrated uncertainty. Confidence distributions for correct versus incorrect Top-1 predictions were compared using Wilcoxon rank-sum tests, with Cohen’s d for effect size and area under the receiver operating characteristic curve (AUC) as a single-number summary of discriminative capacity (correctness as binary outcome, confidence as predictor). Holm correction was applied across strata.

When evaluating multiple studies, Top-k, MRR, and Recall@GT can be aggregated across studies either by pooling all individual queries (*micro*-average) or by computing the metric within each study and averaging them (*macro*-average); macro-averaging preserves the study-to-study variability, while micro-averaging would mask it. We reported micro-averages by default and used macro-averages only for benchmarking against Magneto, where the original paper reported per-study mean across the studies^9^ (**Supplementary Methods**).

## Results

### Performance of *SchemaMapper* with Alias Dictionaries

During *SchemaMapper* algorithm design, we observed that the target attributes expanded with their manually-curated alias dictionary performed better than the no_alias. To ensure the scalability of *SchemaMapper*, we systematically generated target alias dictionaries using different LLM models. Six proprietary alias dictionaries improved Top-1 *SchemaMapper* accuracy over the no_alias baseline by 12.1-21.8 pp (**Supplementary Results** and **Supplementary Table 3**), but the performance among them did not scale with model capability or dictionary size. Although *SchemaMapper* with Opus-4.5-alias produced the highest point estimates (Top-1 = 75.76%, Top-5 = 86.06%, MRR = 0.800, Recall@GT = 0.699, micro-average), it was statistically indistinguishable from five other top-tier dictionaries on Top-1 and Recall@GT (**Supplementary Results**). Within this equivalence set, Claude Haiku 4.5 sat on the Pareto frontier of cost-versus-accuracy, especially with one-time API cost roughly 20× lower than Opus 4.5 and ∼2.8x faster wall-clock time; two Tier-A models (Sonnet 4.5 and Opus 4.7) were each strictly Pareto-dominated by a cheaper same-vendor alternative (**Supplementary Figure 1D** and **Supplementary Table 15**). The open-weight Gemma 3 27B dictionary performed at the no-alias baseline; two additional open-weight models (Gemma 4 26B and Qwen3 32B) failed format compliance and produced no usable dictionary.

*SchemaMapper* performance also varied by pipeline stage (**Figure 2B** and **Supplementary Table 4**): Stage 1 resolved 77 queries at 87.0% Top-1 accuracy, while Stage 3 handled 87 queries at 62.1% Top-1 accuracy. Without aliases, queries previously resolved at Stage 1 fell through to Stage 3 semantic matching, increasing the Stage 3 query pool from 87 to 114 and diluting its within-stage accuracy (Top-1: 62.1% → 38.6%; MRR: 0.692 → 0.495). This demonstrates that the alias dictionary improves performance through two mechanisms: directly, by resolving queries at the most reliable pipeline stage, and indirectly, by reducing the difficulty of the residual semantic-matching pool.

**Figure 1.**
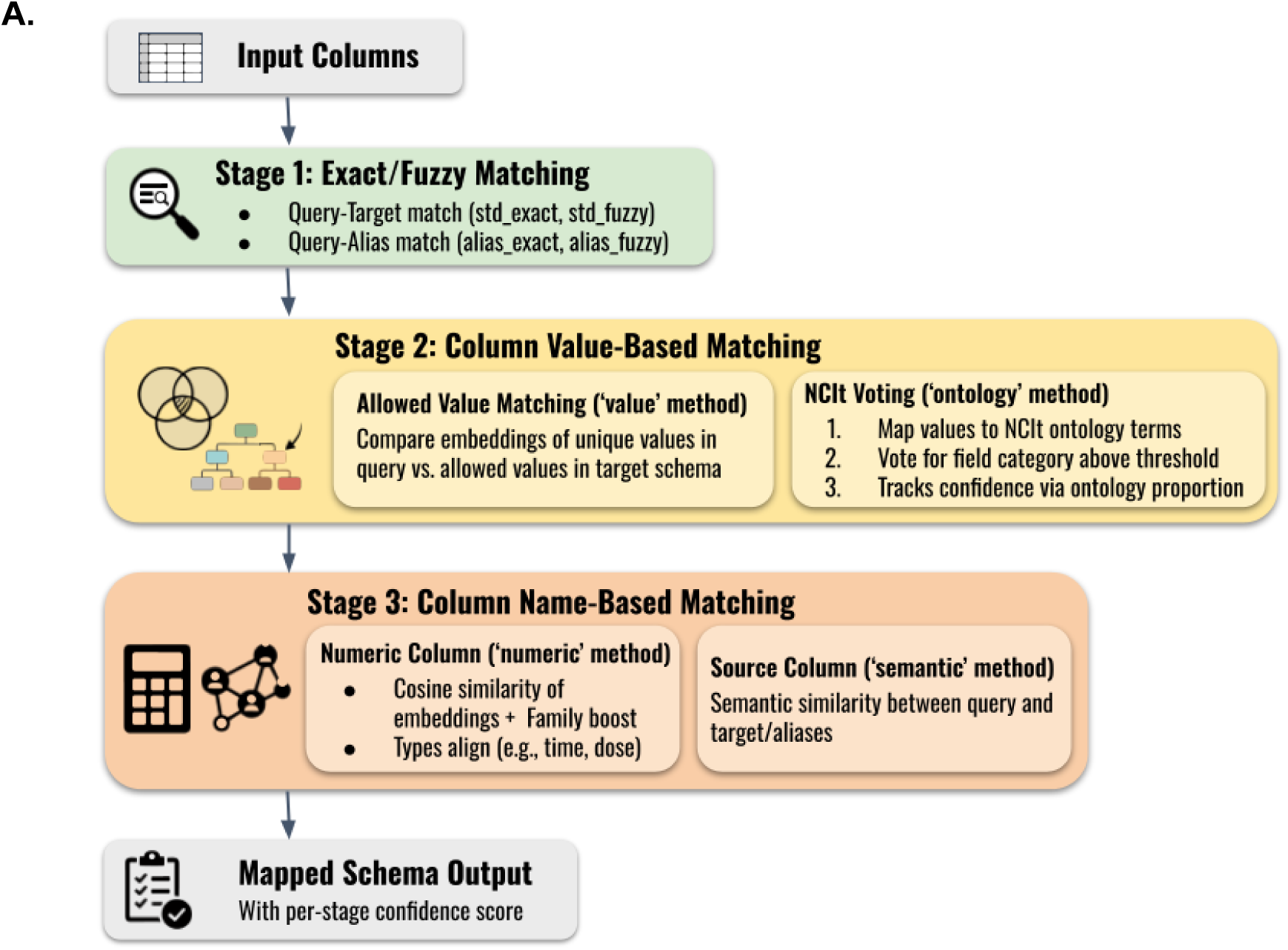
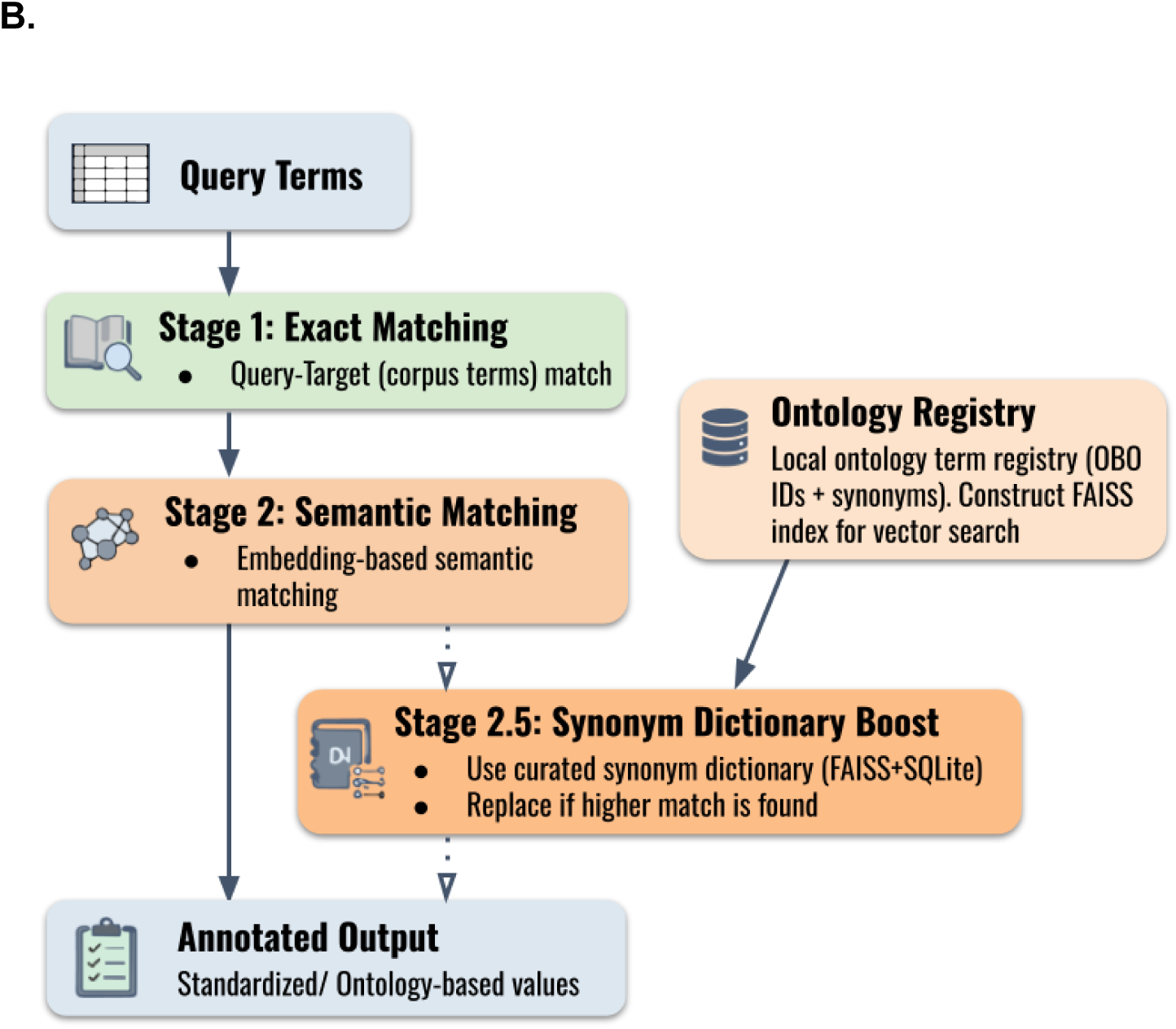
Overview of *MetaHarmonizer*. *MetaHarmonizer* performs metadata harmonization at two levels: attribute-level via *SchemaMapper* and value-level via *OntologyMapper*. Each module cascades queries through stages, applying low-cost lexical methods first and reserving embedding-based methods for unresolved queries. **(A)** *SchemaMapper* aligns input column names to a target schema in three sequential stages. Stage 1 (green) performs exact and fuzzy matching against target names and their aliases. Unresolved queries advance to Stage 2 (yellow), which uses column-content evidence: the ‘value’ method compares embeddings of unique values across source and target columns, while the ‘ontology’ method maps values to ontology terms and votes for a target field when >20% match, tracking confidence via the matched proportion. Remaining queries enter Stage 3 (orange), which compares column-name embeddings: the ‘numeric’ method applies a family-aware boost when types align (e.g., time, dose), and the ‘semantic’ method handles ambiguous headers via target/alias similarity. Each output is annotated with its resolving stage and a per-stage confidence score. **(B)** *OntologyMapper* grounds free-text values to ontology terms in two primary stages, plus a dictionary-boost step. Stage 1 (green) performs exact matching against the corpus terms. If there is no confident match, queries advance to Stage 2 (orange), which retrieves candidates based on cosine similarity between the query and target embeddings, using a FAISS index over a local ontology registry. Low-confidence Stage 2 outputs trigger Stage 2.5, which queries a curated synonym dictionary (FAISS + SQLite) and replaces the candidate if a higher-scoring match is found. The outputs are standardized, ontology-grounded values.

**Figure 2.**
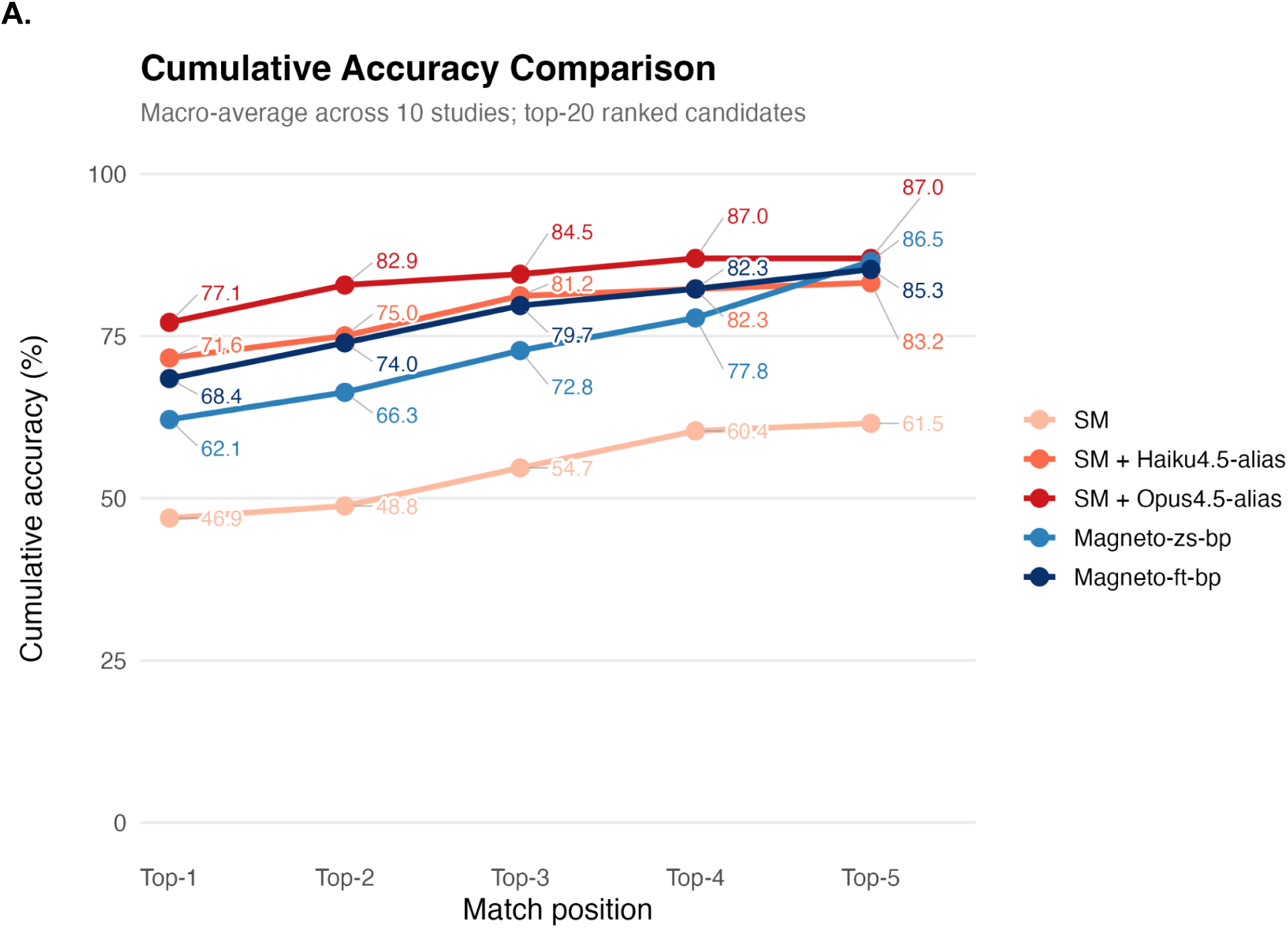
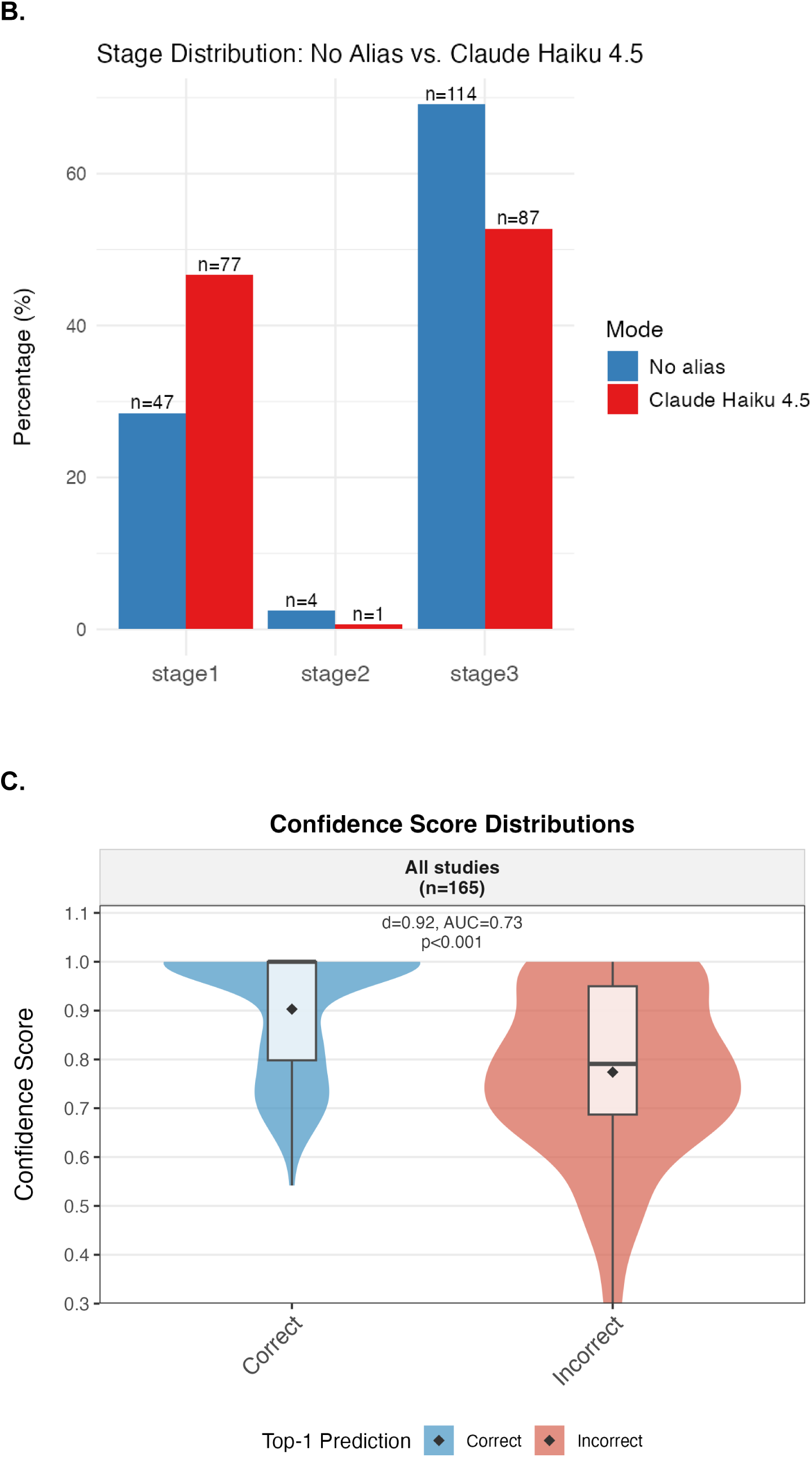
Performance of *SchemaMapper* on the GDC schema mapping benchmark. We benchmarked *SchemaMapper* against 165 ground-truth column mappings from the GDC schema matching data. **(A)** Top-*k* accuracy (k = 1-5) for five configurations: Magneto zero-shot bipartite reranker (Magneto-zs-bp, blue), Magneto fine-tuned bipartite reranker (Magneto-ft-bp, navy), *SchemaMapper* without an alias dictionary (SM, peach), and *SchemaMapper* with a Haiku-4.5-alias (SM + Haiku4.5-alias, orange) and an Opus-4.5-alias (SM + Opus4.5-alias, red) **(B)** Stage distribution of all queries with and without the Haiku-4.5 alias dictionary. The dictionary expands Stage 1 coverage from 47 to 77 queries and reduces Stage 3 coverage from 114 to 87 queries; Stage 2 query counts are barely changed. The shift indicates that the alias dictionary resolves additional queries through low-cost lexical matching that would otherwise require semantic embeddings. **(C)** Confidence calibration for the SM + Haiku4.5-alias. Paired violin and box plots compare the score distribution of correct (blue, n = 121) and incorrect (red, n = 44) Top-1 predictions. Correct predictions concentrate near 1.0 (mean = 0.90, median = 1.00), whereas incorrect predictions are more dispersed (mean = 0.77, median = 0.79), yielding strong separation (Cohen’s *d* = 0.92, AUC = 0.73, Wilcoxon-p < 0.001). The score is therefore well calibrated for triaging predictions to human review. Panels B and C used SM with Haiku4.5-alias because it is the recommended model for deployment. SM = *SchemaMapper*.

### *SchemaMapper* Benchmarking

We benchmarked *SchemaMapper* with Haiku-4.5-alias, the recommended deployment option, against four Magneto configurations on the GDC schema-matching benchmark (**Table 1** and **Figure 2A**). *SchemaMapper* outperformed both reproducible Magneto bipartite variants (zs-bp and ft-bp) on Top-1, MRR, and per-query Recall@GT: Top-1 accuracy (71.6% vs. 62.1% and 68.4%), MRR (0.763 vs. 0.731 and 0.757), and per-query Recall@GT (0.65 vs. 0.537 and 0.60), with the difference against zs-bp reaching statistical significance (p = 0.033, 0.042, and 0.013). With the best-performing alias dictionary (Opus-4.5-alias), *SchemaMapper* was significantly better than both Magneto bipartite variants on Top-1, MRR, and per-query Recall@GT (p = 0.032, 0.03, and 0.014 for zs-bp; p = 0.03, 0.033, and 0.032 for ft-bp); it is also statistically indistinguishable from Magneto LLM-reranked variants on MRR (per-study paired Wilcoxon-p = 0.93 (zs-llm) and 0.43 (ft-llm)). Per-query Recall@GT complements MRR for this benchmark, where 40% of queries (66/165) have multi-target ground truth. *SchemaMapper*’s confidence scores were well calibrated for distinguishing correct from incorrect predictions (**Figure 2C**): mean confidence was 0.903 for correct predictions (n = 121) and 0.774 for incorrect predictions (n = 44), with Cohen’s d = 0.92 and AUC = 0.73.

**Table 1.** Benchmark comparison of *SchemaMapper* and *Magneto* on the GDC datasets. Ten CPTAC data tables were mapped to the 736-column GDC target schema, and performance was evaluated using ground-truth matches for 165 source columns created through expert curation. We compared the no-alias SM baseline, SM with three LLM-generated alias dictionaries (the best vendor-diverse option; see Supplementary Results), and four Magneto variants. Recall@GT is reported under two non-interchangeable definitions, *per-query* and *global*, which differ in whether they penalize cross-query score miscalibration; the two columns should not be read as adjacent values (see Supplementary Methods for definitions and the conditions under which each is computable). *SchemaMapper* produces a sparse candidate list and cannot support the global Recall@GT definition, and Magneto-llm rows do not include the per-query rank distributions needed to compute Top-1 and Top-5. All cells are *macro-averaged* across studies (per-study metric first, then unweighted mean over the ten studies). Asterisked (*) Magneto LLM variants are paper-reported (Liu et al.^1^ Table 6, MPNet rows), not reproduced locally. Daggered (†) Magneto bipartite variants were reproduced locally, matching the source paper within ±0.005 on both MRR and global Recall@GT. “—” represents a metric that is unavailable in the source data. SM = *SchemaMapper*; zs = zero-shot; ft = fine-tuned; bp = bipartite-reranker; llm = LLM-reranker.

| Method | Training | MRR (macro) | Top-1 (%) | Top-5 (%) | Recall@GT (per-query) | Recall@GT (global) |
| --- | --- | --- | --- | --- | --- | --- |
| SM (no alias) | Zero-shot | 0.521 | 47 | 61.5 | 0.425 |  |
| SM + Opus 4.5 alias | Zero-shot | 0.811 | 77.1 | 87 | 0.685 | — |
| SM + Haiku 4.5 alias | Zero-shot | 0.763 | 71.6 | 83.2 | 0.65 | — |
| SM + Gemini 2.5 Pro alias | Zero-shot | 0.76 | 72.8 | 80.9 | 0.659 | — |
| Magneto-zs-bp † | Zero-shot | 0.731 | 62.1 | 86.5 | 0.537 | 0.373 |
| Magneto-ft-bp † | Fine-tuned | 0.757 | 68.4 | 85.3 | 0.601 | 0.433 |
| Magneto-zs-llm * | Zero-shot | 0.808 | — | — | — | 0.43 |
| Magneto-ft-llm * | Fine-tuned | 0.846 | — | — | — | 0.537 |

### *OntologyMapper* Benchmarking

*OntologyMapper* consistently outperformed *text2term* across all four benchmarks on both Top-1 and MRR (**Figure 3A** and **Table 2**). The largest margins were on Biomappings-EFO (Top-1 95.5% vs. 79.1% (Δ = +16.4pp); MRR 0.969 vs. 0.848), followed by OLS-EFO full (Top-1 89.1% vs. 79.2% (Δ = +9.9pp); MRR 0.903 vs. 0.813). On UKBB-EFO, *OntologyMapper* reached 77.9% Top-1 vs. *text2term*’s 71.6% (MRR 0.826 vs. 0.765). The narrowest gap was on OLS-EFO disease (Top-1 95.2% vs. 93% (Δ = +2.2pp); MRR 0.963 vs. 0.952), the easiest of the four benchmarks, where both tools approach the ceiling set by the curated ground truth.

**Figure 3.**
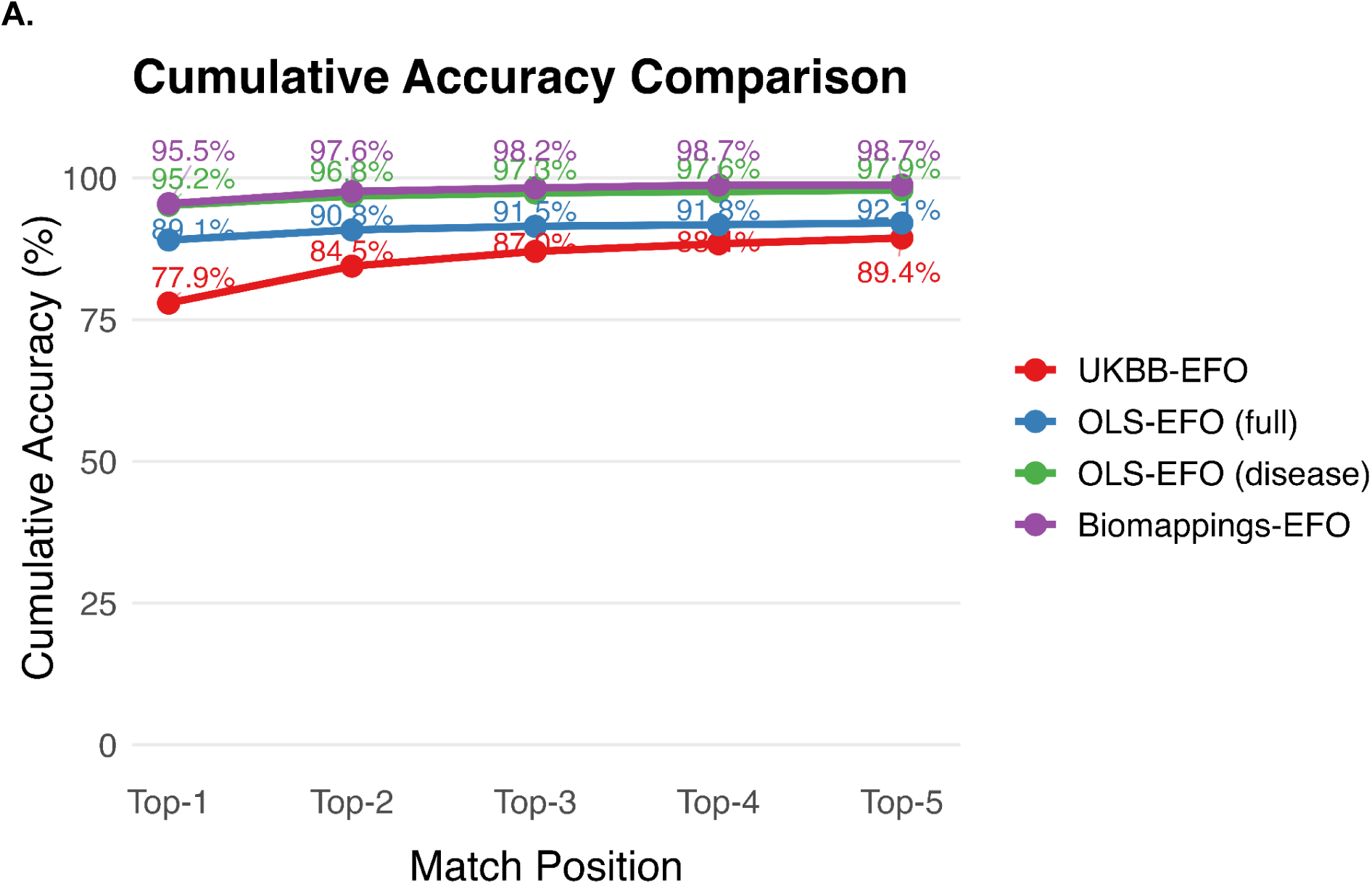
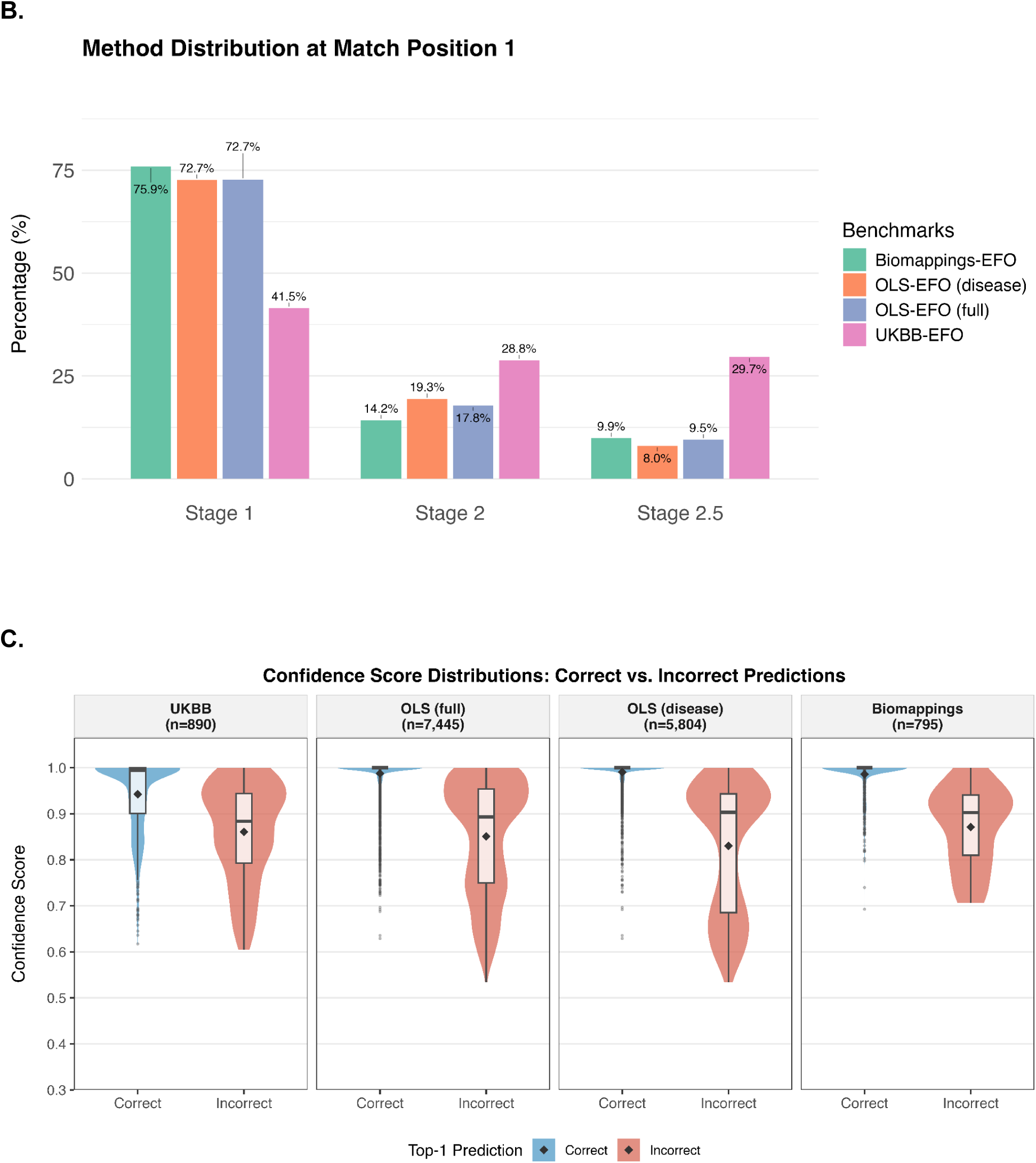
Performance of *OntologyMapper* across diverse benchmarking datasets. We evaluated *OntologyMapper* on four benchmarks: Biomappings-EFO, OLS-EFO (disease subset), OLS-EFO (full), and UKBB-EFO. **(A)** Top-k cumulative accuracy (k = 1-5) on each benchmark. The gap between Top-1 and Top-5 accuracy is the largest for UKBB-EFO. **(B)** Composition of correct Top-1 predictions by resolving stage, stratified by benchmark. Stage 1 resolved the majority of queries on benchmarks with formal ontology labels as input (Biomappings-EFO, OLS-EFO disease, and OLS-EFO full), whereas UKBB-EFO depends more heavily on the embedding-based stages (Stages 2 and 2.5), consistent with the greater lexical divergence of UK Biobank phenotype descriptions from EFO labels. **(C)** Confidence score distributions for correct (blue) and incorrect (red) Top-1 predictions, faceted by benchmark (*n* shown above each panel). Paired violin and box plots show the full density of scores; embedded box plots indicate the median (horizontal line), interquartile range (box), and 1.5× IQR whiskers; diamond markers indicate the mean. The prominent density spike at 1.0 among correct predictions reflects Stage 1 terms resolved with perfect confidence. The consistent separation across all benchmarks indicates that confidence thresholding can effectively triage predictions for human review.

**Table 2.**
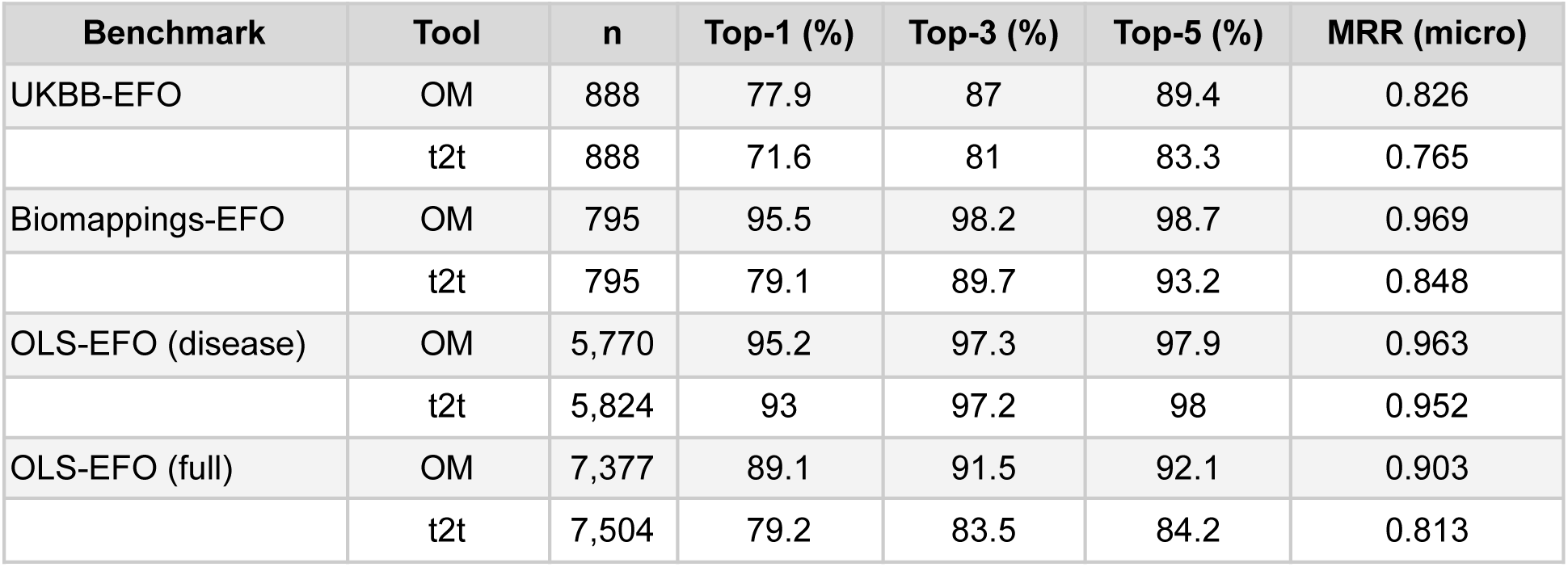
Benchmark comparison of *OntologyMapper* and *text2term* across four EFO mapping datasets. Ontology mapping performance was evaluated on four ground-truth benchmarks of increasing scale and heterogeneity. The target corpus includes the 12-ontology EFO selection (33,230 terms for OM and 33,659 terms for t2t). For each benchmark, *n* indicates the number of query terms with at least one ground-truth target. Small differences in *n* between tools reflect tool-specific handling of queries that yield no candidates within the configured search space. *OntologyMapper* outperforms *text2term* across all four benchmarks and metrics, with the largest absolute gains observed on Biomappings-EFO and the smallest on OLS-EFO (disease), where both tools approach ceiling performance. We used the Top-5 retrievals and the micro-average for this evaluation. OM = *OntologyMapper*; t2t = *text2term*.

The multi-stage pipeline contributed differentially across benchmarks (**Supplementary Table 5**). Stage 1 resolved 32.4–72.5% of queries (median ∼67%), accounting for 41.5-75.9% of correct Top-1 predictions (**Figure 3B**). Among queries routed to Stage 2, Top-1 accuracy ranged 70.4–88.6% at Top-1, and Stage 2.5 resolved queries unmatched by either lexical or semantic match at 59.2–81.5% Top-1.

Confidence scores showed statistically significant separation between correct and incorrect Top-1 predictions across all four benchmarks (Wilcoxon rank-sum test, all p < 2 × 10⁻¹⁶ after Holm correction; **Figure 3C** and **Supplementary Table 6**). Effect sizes were large, with Cohen’s d ranging from 0.954 (UKBB-EFO) to 3.89 (OLS-EFO disease). The AUC for confidence-based discrimination of correct versus incorrect predictions ranged from 0.766 (UKBB-EFO) to 0.935 (OLS-EFO disease), indicating that confidence scores provide a useful signal for identifying predictions that may require manual review. Mean confidence was substantially higher for correct than for incorrect predictions across all benchmarks: 0.94 vs. 0.86 (UKBB-EFO), 0.99 vs. 0.85 (OLS-EFO full), 0.99 vs. 0.83 (OLS-EFO disease), and 0.99 vs. 0.87 (Biomappings-EFO). The lower AUC on UKBB-EFO is consistent with the greater lexical diversity of UK Biobank phenotype descriptions, which span a wider range of difficulty levels, compressing the confidence distribution.

### Runtime Analysis

To evaluate the computational efficiency, we profiled *SchemaMapper* and *OntologyMapper* across their respective benchmark tasks, measuring end-to-end pipeline time and scalability with respect to input size.

The GDC benchmark datasets for *SchemaMapper*+Haiku-4.5-alias include 10 studies ranging 5-29 source columns each. The multi-stage cascade triages columns by difficulty: 46.7% of columns (77/165) were resolved at Stage 1 at negligible cost (<0.001s per column, mean ≈ 55µs), while the remaining 88 (53.3%) advanced to Stages 2/3, with 87/88 terminating at Stage 3. Stage 2 carried the highest per-query latency (mean 0.39s) and dominated cumulative compute (29.7s of 35.7s total wall-clock), while Stage 3 added only 0.08s per query on average for 736-way semantic search. This concentration of compute at Stage 2 reflects the cost of pairwise alias-and-value comparison; Stage 3’s amortized FAISS search is comparatively cheap. End-to-end pipeline time ranged from 0.52s (Krug: 5 columns) to 12.7s (Cao: 29 columns). Total Stage 3 time scaled approximately linearly with the number of columns reaching that stage (Pearson r = 0.868), consistent with the expected linear cost of FAISS search; the full pipeline runtime showed a similar pattern (Pearson r = 0.797) (**Figure 4A**).

**Figure 4.**
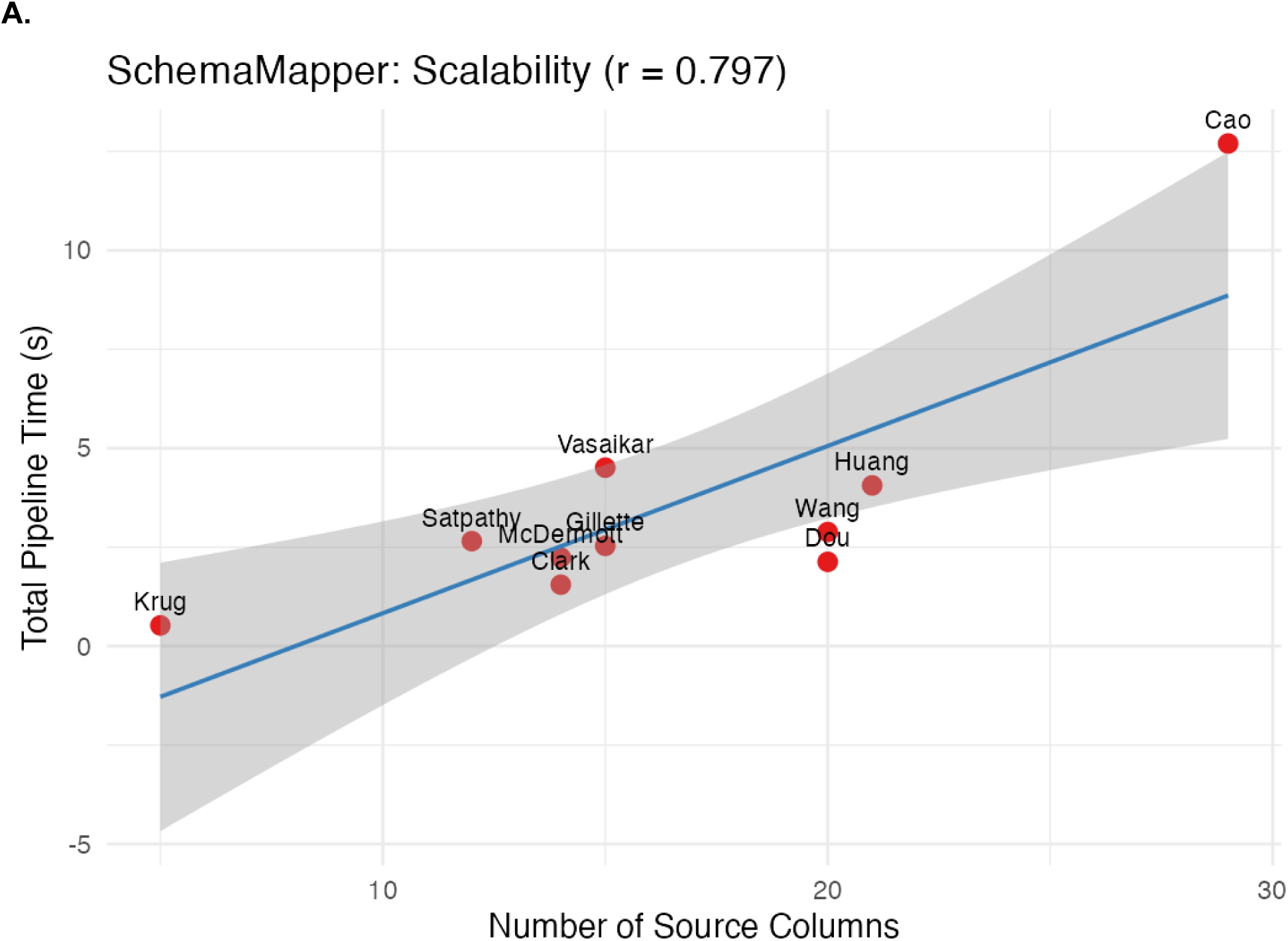
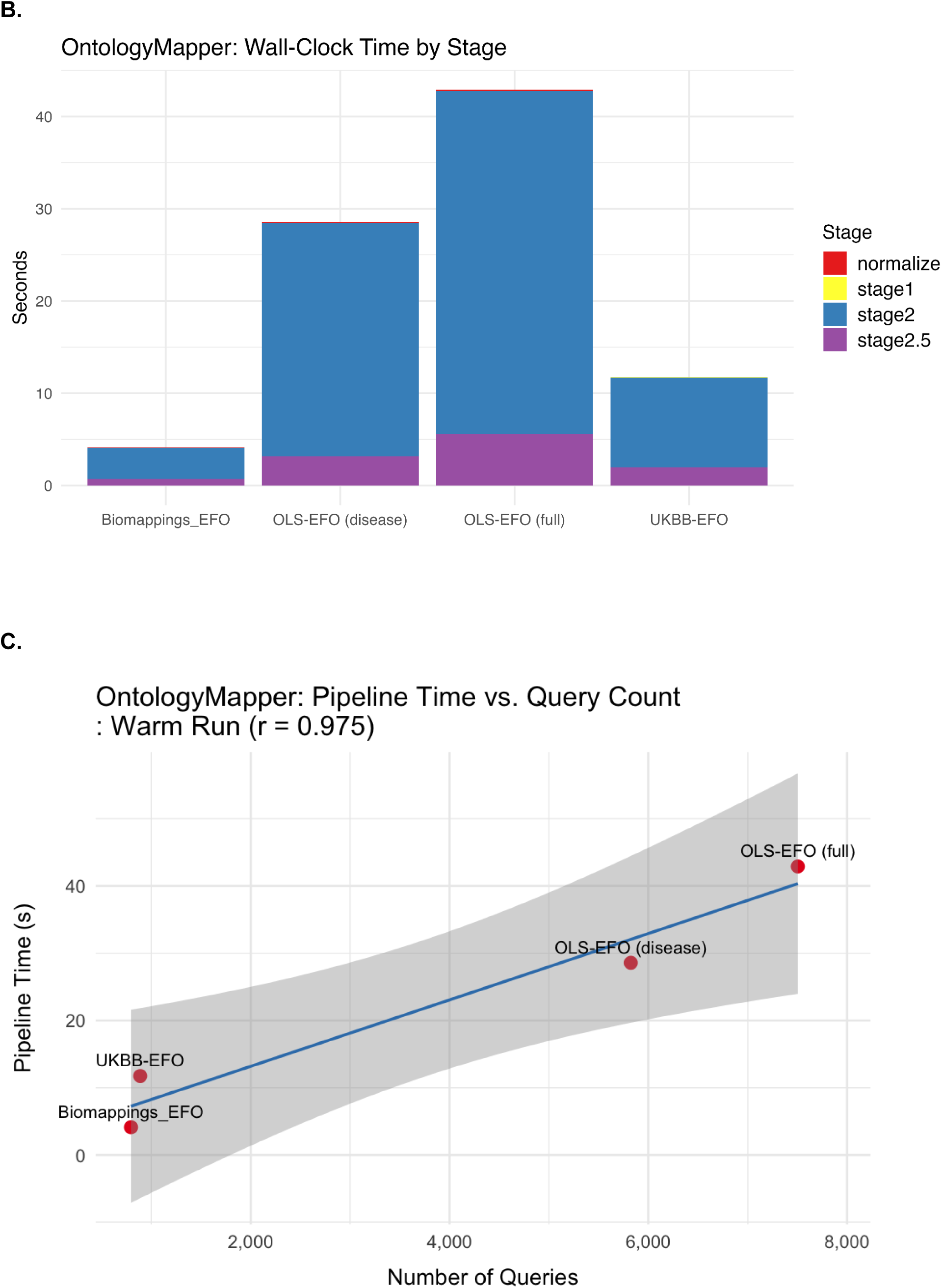
Runtime analysis of *MetaHarmonizer*. We profiled the runtime of *MetaHarmonizer* modules on representative benchmarks. The target corpus contains 736 GDC columns for *SchemaMapper* and 33,230 EFO terms for *OntologyMapper*. **(A)** *SchemaMapper* runtime (with the Haiku-4.5-generated alias dictionary) scales approximately linearly with the number of source columns. Each point represents one of the 10 CPTAC benchmark studies (labeled by first author) and plots total pipeline time (seconds) against the number of source columns. The blue line is a linear regression fit with 95% confidence interval (grey shading) (Pearson *r* = 0.797). **(B)** *OntologyMapper* wall-clock time broken down by stage, on each of the four EFO benchmarks. *OntologyMapper* Stages 2 and 2.5 together account for nearly all active pipeline time; Stage 1 contributes negligibly. **(C)** *OntologyMapper* warm-run scaling across the four EFO benchmarks. “Warm” denotes a run in which the FAISS index is already loaded into memory and embeddings are cached, so per-query cost reflects steady-state performance rather than initialization. Pipeline time scales approximately linearly with query count (Pearson *r* = 0.975).

We profiled *OntologyMapper* across four EFO benchmark datasets spanning a ∼10-fold range of query sizes, all mapped against a 33,230-term EFO corpus (**Figure 4B** and **C**). Cold (first-time) runs were dominated by initialization, taking ∼1,200s (∼20min) per benchmark regardless of query count, because the corpus pass (embedding 33,230 terms, ∼7.5 min) and the synonym pass (embedding ∼142,645 texts, ∼11min). This cost is incurred only once per *(model, corpus-content)* pair and is amortized across subsequent runs through the warm cache (**Methods**). On subsequent warm runs, the initialization cost drops to under 0.4s, reducing the total runtime from average ∼21.5min to ∼22s per benchmark (**Supplementary Table 7**): Biomappings-EFO was fastest (4.45s) because 72.5% of queries were resolved at Stage 1. OLS-EFO-full was the slowest (43.2s), reflecting its query volume, combined with k=5 FAISS search and alias rerank, on a ∼36% non-exact residual. For comparison, *text2term* ran at a consistent ∼15s per benchmark from local corpus, regardless of query count (**Supplementary Table 8**).

### Contamination-controlled evaluation reveals memorization as a confounder in LLM-only schema matching

On the GDC schema-matching benchmark, five frontier LLMs (Claude Haiku 4.5, Sonnet 4.5, and Opus 4.5; Gemini 2.5 Flash and Gemini 2.5 Pro) reached 83.0–89.7% Top-1 accuracy, exceeding *SchemaMapper* with a corresponding alias dictionary by +10.3 to +23.6 pp (matched-model paired comparison; Wilcoxon p < 1 × 10⁻⁵; **Supplementary Table 10**). However, 131 of 165 queries (79.4%) were resolved correctly by all five models simultaneously. Such cross-family agreement is uncommon under independent reasoning and is suggestive of a shared exposure to GDC documentation in pretraining data rather than transferable matching capability^32,33^.

Direct memorization probes (E4 with probes P1-P3, defined in **Supplementary Methods**) test for memorized GDC structure independent of the matching task. (**Supplementary Table 11**). Three out of five frontier models recovered 80–100% of GDC identifiers from zero-schema context (P2), with matched performance across the Anthropic and Google model families, ruling out a single-lab data-curation artifact; near-zero exact recovery on a positional completion probe (P3: Gemini 2.5 Pro 7%, all others 0%) indicates that this knowledge is indexed by clinical concept rather than stored as a verbatim copy of the 736-column list. For the GDC benchmark test, target-vocabulary rewriting (E3) erased the LLM-only advantage (**Supplementary Methods** and **Supplementary Table 12**). All 736 GDC target identifiers were renamed to semantically equivalent non-GDC strings; this collapsed LLM-only Top-1 by 12.1–23.6 pp while *SchemaMapper* was essentially unchanged (−1.8 to +3.0 pp) (**Supplementary Table 12**). Critically, the paired LLM-only vs. *SchemaMapper* gap collapsed from -10.3 pp at baseline to +7.3 pp under target rename for Haiku 4.5, and from -23.6 pp to +2.4 pp for Sonnet 4.5; across all five matched pairs, *SchemaMapper* paired with an LLM-generated alias dictionary now outperforms the matched LLM-only counterpart by +2.4 to +7.3 pp (**Figure 5**).

**Figure 5.**
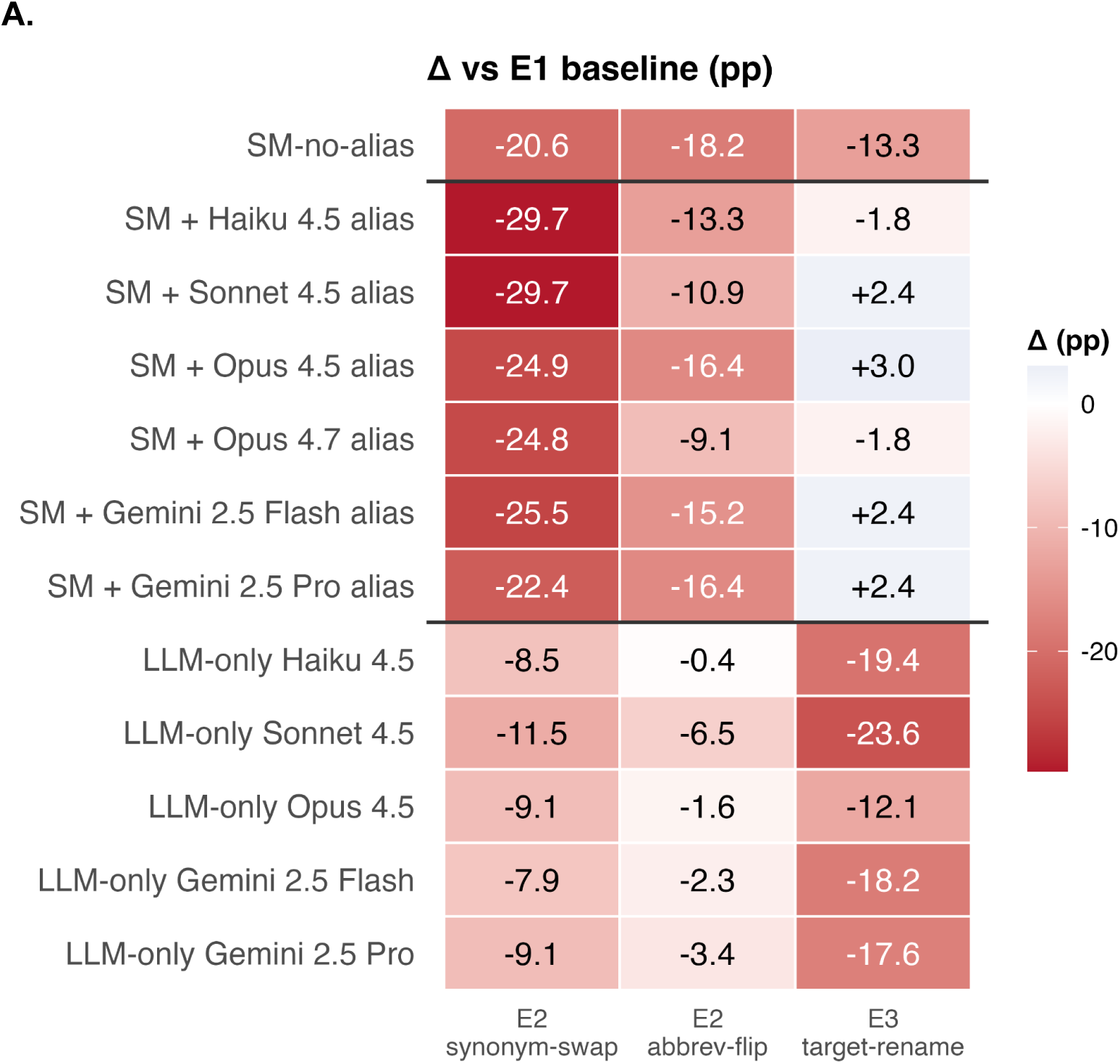
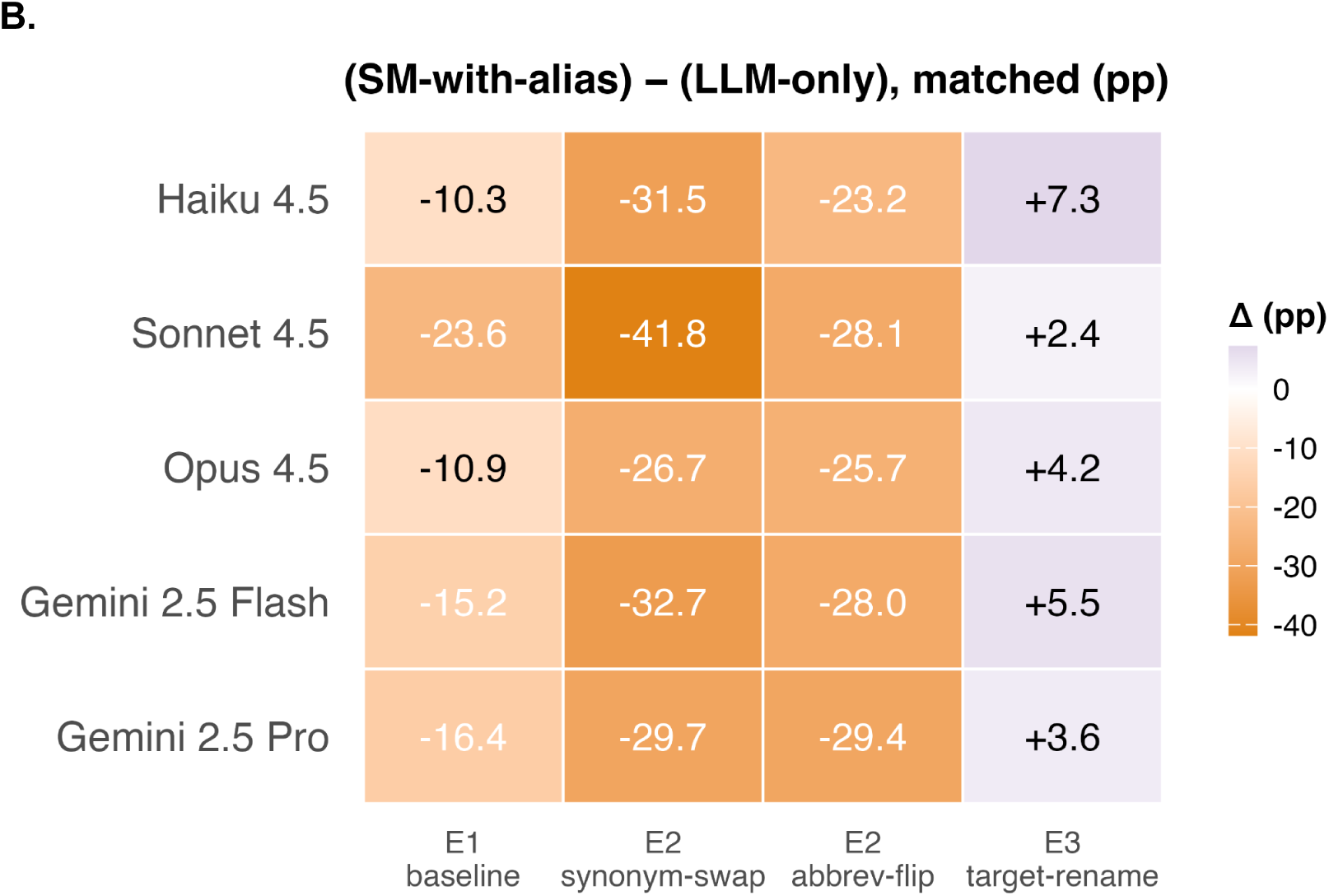
LLM-contamination on schema mapping. **(A)** Per-configuration change in Top-1 accuracy under each of the three perturbation conditions, computed as ‘*condition − E1-baseline*’ (Δ). E2 comprises two source-side variants reported separately: synonym-swap and abbreviation-flip. Rows are the 12 configurations grouped into three families separated by horizontal lines: SM with no alias (*SM-no-alias*), SM augmented with LLM-generated aliases (six variants), and LLM-only baselines (five variants; Claude Opus 4.7 was excluded because it does not accept temperature=0). Cell color encodes the signed Δ (red = accuracy loss, blue = accuracy gain). **(B)** Difference in Top-1 accuracy between SM with LLM-alias dictionaries and the LLM-only baseline using the *same* underlying model, computed as ‘*SM-with-alias − LLM-only*’, shown for each of the four conditions (E1 baseline plus three perturbations). Rows are the five matched-model pairs; Opus 4.7 has no LLM-only counterpart in the table and is omitted. Cell color encodes the signed gap (orange = *SM-with-alias* is worse than the LLM-only baseline, purple = *SM-with-alias* is better). When the GDC target labels are renamed (E3), all five *SM-with-alias* configurations overtake their LLM-only counterparts. SM = *SchemaMapper*; cell text reports values as *±pp*.

These contamination-controlled evaluations provide direct evidence that GDC concept-to-identifier knowledge is present in both pretraining corpora and confirm the mechanism inferred from the E3 target-rename collapse. LLM-only retained one genuine advantage: robustness to source-column paraphrase, where LLM performance degrades more gracefully than *SchemaMapper*’s retrieval (**Supplementary Results**).

### *OntologyMapper* outperforms LLM-only baselines where memorization is not a shortcut

We evaluated LLM-only performance on the most challenging EFO benchmark datasets, UKBB-EFO. Because the 33,230-term merged corpus exceeds Claude Haiku 4.5’s input token limit, this evaluation used a 17,638-term EFO-native corpus; *OntologyMapper* was re-run on the same 17,638-term corpus for a matched comparison. The LLM-only baseline fell ∼19-28 pp short of *OntologyMapper* under both zero-shot and open-book prompting (**Supplementary Table 13**). Closed-book zero-shot prompting reached 54.9% Top-1 (MRR = 0.584) with a 34.5% hallucination rate; injecting the full 17,638-label EFO corpus as a cached system block lifted accuracy to 63.7% Top-1 (MRR = 0.676) and reduced hallucination to 6.0%, but still underperformed *OntologyMapper* (83.0% Top-1, MRR = 0.887; 17,638-term corpus) at a cost of ∼$15 (USD) per benchmark run (input: ∼322k cache-write + ∼143.2M cache-read + ∼35k uncached; output: ∼129k tokens, totaling ∼$15.40 (USD) at Claude Haiku 4.5 (claude-haiku-4-5-20251001) standard real-time API rates as of 2026-04-20 (cache-read alone accounts for ∼$14.32)). Per-query agreement analysis between *OntologyMapper* and open-book LLM-only showed *OntologyMapper* was uniquely correct on 24.4% of queries and LLM-only was uniquely correct on only 5.2%, with the remainder either both correct (58.8%) or both wrong (11.6%), indicating *OntologyMapper* encompasses most of the LLM’s capability while contributing substantial unique coverage.

## Discussion

### Reported accuracies of LLM-based metadata harmonization on public biomedical benchmarks can substantially overstate transferable capability

Because this pattern is observed across two independent pretraining corpora (Claude and Gemini), it is unlikely to be a single-lab curation artifact and instead points to widespread exposure of public schema documentation in pretraining data. The implication extends well beyond *MetaHarmonizer*: any LLM-augmented bioinformatics evaluation built on widely-documented public reference resources (e.g., GDC, GTEx variable dictionaries, TCGA harmonized schemas) should be interpreted as an upper bound that may not transfer to novel or proprietary schemas absent from pretraining. The contamination battery (**Supplementary Methods**) is task-agnostic and can be applied wherever an LLM baseline is reported against a public benchmark. This kind of probing may be a useful addition to LLM-augmented benchmarks, in the same way that train-test leakage checks are now standard for supervised learning evaluations.

This finding reframes the methodological question for biomedical metadata harmonization; not whether LLMs can match documented schemas they may have memorized, but how to build harmonization systems that remain reliable on novel schemas, in offline settings, and under the FAIR-compliant data infrastructure. *MetaHarmonizer* is one response to that question: a multi-staged retrieval architecture that uses LLMs only as bounded components within a deterministic and auditable pipeline, achieving competitive accuracy with grounded outputs, calibrated confidence, and stable performance under contamination probes.

### LLMs as components rather than as solutions

*SchemaMapper* uses LLMs for one targeted role – alias dictionary generation as one-time preprocessing – while grounding all outputs in a pre-defined schema. Its performance was stable across the contamination probing conditions because the core matching algorithm depends on embedding similarity to target column names rather than memorized schema knowledge. This robustness is from the architectural choice: contamination affects the alias dictionary’s coverage but not the deterministic matching that follows.

### Aliasing matters, but the choice of the LLM model for alias generation mostly does not

A prevalent assumption in LLM-augmented bioinformatics is that a more capable model uniformly improves output quality. We find a weaker pattern for embedding-friendly synonym generation: six top-tier proprietary models generated alias dictionaries whose downstream *SchemaMapper* performance was statistically indistinguishable on Top-1 accuracy after multiple-comparison correction, despite spanning roughly a 50-fold range in API cost. The failure mode of larger models is not incorrect aliases but too many correct-but-off-task ones, such as pipeline tool names and value-like strings that are plausible surface forms yet poor semantic neighbors of the actual queries. Because *SchemaMapper* Stage 3 ranks aliases by cosine similarity, these embedding-unfriendly aliases pull unrelated queries toward the wrong field or dilute rank by occupying independent embedding slots. The practical consequence is that Haiku-4.5 is the Pareto-optimal default for *SchemaMapper* alias dictionary generation (**Supplementary Figure 1** and **Supplementary Table 15**). The implication generalizes beyond *MetaHarmonizer*: in any retrieval pipeline that consumes LLM-generated text via dense embeddings, model selection should be aligned with the downstream consumer, not general capability benchmarks. This suggests that within the equivalence set, users can pick on cost, latency, vendor, or compliance option without compromising accuracy.

Beyond accuracy, the LLM-generated alias dictionary requires only target column names as input, making adaptation to new schemas a matter of providing column names and, optionally, modified prompt templates. Direct cross-schema validation remains future work, but the *source-agnostic generation procedure* provides easy adaptation to new domains.

### *SchemaMapper* recovers more plausible candidates for human review

With the LLM-generated alias dictionary, *SchemaMapper* achieved the highest per-query Recall@GT among the tested, including Magneto’s fine-tuned bipartite variant. This advantage reflects a benefit of our multi-stage cascade design: it draws candidates from structurally different matching strategies, each with distinct biases, and naturally surfaces multiple acceptable targets within its candidate list. In real-world schema harmonization where one-to-many mappings are common, curating a diverse high-quality candidate set for human review is often more valuable than rank-1 precision alone. Magneto’s fine-tuned LLM-reranker on MRR is statistically indistinguishable from *SchemaMapper*+Opus-4.5-alias, and the bipartite reranker configurations are beaten by *SchemaMapper*+Opus-4.5-alias and statistically indistinguishable from *SchemaMapper*+Haiku-4.5-alias. Thus, the two systems can occupy complementary points on the precision–recall frontier: auto-fill workflows benefit from Magneto’s LLM-reranker, while human-in-the-loop review benefits from *SchemaMapper*.

### *OntologyMapper*’s accuracy advantage stems from multi-stage semantic matching

*OntologyMapper* achieved Top-1 accuracy of 77.9–95.5% and MRR of 0.83–0.97 across four EFO benchmarks (on a 33,230-term corpus, micro-average), consistently outperforming *text2term*. The performance gap derives primarily from semantic matching stages; SapBERT was pre-trained on millions of UMLS synonym pairs and place semantically equivalent terms close together regardless of lexical overlap, while *text2term*’s TF-IDF backbone is sensitive to surface-form variation. Performance variation across benchmarks is interpretable from source-term properties: UKBB-EFO yielded the lowest accuracy because UK Biobank phenotype descriptions are often informal or use lay terminology (e.g., “blood clot in the lung” vs. “pulmonary embolism”), reducing Stage 1 exact-match resolution. The OLS-EFO disease subset outperformed the full benchmark because disease ontology terms have more extensive synonym sets and more consistent naming conventions. The two tools occupy complementary positions: *text2term* as a lightweight, broadly accessible solution well-suited to lexically aligned tasks; *OntologyMapper* as a higher-accuracy option for semantically challenging mappings, at the cost of transformer embeddings.

### Cascade design makes the system tractable

Each module’s “easy-first” design resolves simple cases before invoking expensive methods. *OntologyMapper*’s Stage 1 provides a high-confidence, zero-error pathway for terms with direct lexical matches, resolving 32.4–72.5% of queries before embedding-based stages are invoked. Stage 2 handles the bulk of semantically complex matching, while Stage 2.5 provides a safety net through expanded synonym search. Beyond accuracy, this organization yields predictable runtime: *SchemaMapper* remains under 30 sec even for the largest CPTAC studies, and *OntologyMapper*’s per-query marginal cost of 0.006s suggests the pipeline accommodates tens of thousands of terms without bottlenecks. *OntologyMapper*’s cold/warm asymmetry reflects one-time embedding passes cached per corpus, a cost model well-suited to typical usage where a laboratory or consortium maps multiple studies against the same target ontology over time.

### Calibrated confidence enables practical triage

Both modules produce confidence scores with strong separation between correct and incorrect predictions, and this separation holds consistently across studies and benchmarks. In production workflows where curators must validate predictions, well-calibrated confidence is often more useful than marginal accuracy gains: predictions above a chosen threshold can be auto-accepted while those below are flagged for review. This thresholding capability is a direct consequence of staged retrieval – each stage’s score reflects a specific matching mechanism with bounded failure modes – and is harder to obtain from end-to-end approaches that produce a single output.

### Limitations

Several limitations should be acknowledged. The contamination battery establishes that LLM-only advantages on the GDC benchmark disappear upon target renaming, but the magnitude and form of contamination effects on other public schemas have not been characterized. Benchmarks span two domains – cancer genomics and disease ontology – and broader coverage would strengthen claims of generalizability. Each module was evaluated against a single primary comparator; the broader landscape of schema matchers (e.g., Valentine^31^, COMA^34^) and ontology mappers (e.g., ZOOMA^35^) was not exhaustively benchmarked. Finally, while *MetaHarmonizer* is designed to be domain-extensible (i.e., target schemas and ontology corpora are user-supplied rather than hard-coded), empirical validation outside the benchmarked domains remains future work.

### Future directions

Applying the contamination protocol systematically across other LLM-augmented bioinformatics benchmarks built on public reference resources is a natural next step. For *MetaHarmonizer* specifically, applying the framework to additional repositories and data types would both extend its practical utility and stress-test its coverage on more heterogeneous real-world inputs. More broadly, harmonization is one component of a larger curation workflow, and integration with complementary tools (e.g., those that extract structured metadata from unstructured sources) could position *MetaHarmonizer* within an end-to-end metadata pipeline. Each of these directions builds on, rather than reorganizes, the *MetaHarmonizer* foundation established here.

## Conclusions

*MetaHarmonizer* demonstrates that staged retrieval with bounded LLM components is a practical architecture for biomedical metadata harmonization at scale. *MetaHarmonizer* is locally deployable and deterministic, and its calibrated confidence scores translate accuracy into actionable triage. As biomedical data repositories continue to expand and cross-study analyses become routine, harmonization frameworks will need to scale not only computationally but also in the trust they earn from curators and downstream users. *MetaHarmonizer* is built for both.

## Data Availability

Benchmark datasets used in this study are available from the corresponding source repositories: https://github.com/VIDA-NYU/magneto-matcher for GDC schema mapping, and https://github.com/rsgoncalves/text2term-evaluation for EFO ontology mapping.

## Code Availability

The *MetaHarmonizer* Python package is available in this GitHub repository: https://github.com/shbrief/MetaHarmonizer.

## Acknowledgements

We thank the cBioPortal community at Memorial Sloan Kettering Cancer Center for serving as our Google Summer of Code host organization. The content is solely the responsibility of the authors and does not necessarily represent the official views of the National Institutes of Health.

## Author Contributions

S.O. and S.D. conceptualized the study. S.O., S.D., C.L., A.D., and M.W. designed the methodology. S.O., C.L., and A.D. performed the investigation, developed the software, and wrote the original draft. S.O. and C.L. conducted the formal analysis and validated the results. S.O., K.G.-P., and K.L. curated the data and provided resources. S.O. created the visualizations. S.O. and S.D. supervised the project. S.O., S.D., and I.d.B. administered the project and acquired funding. All authors reviewed and edited the manuscript.

## Supplementary Methods

### Schema-matching evaluation: metric definitions and aggregation choices

Cross-method comparison on the GDC schema-matching benchmark involves several closely related but non-equivalent metrics, two distinct aggregation schemes, and one cascade-specific implementation property that affects reported numbers. We document each below.

#### Per-query metrics

For each source column *c*, each method returns a ranked list of candidate target columns 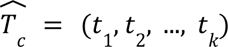, truncated at *K* predictions. Let G*c* denote the set of ground-truth targets for *c* with |*G_c_*| = *n_c_*. We compute three per-query metrics from this ranked list.

- *Mean Reciprocal Rank (MRR).* Following Liu et al.^1^, the per-column reciprocal rank is *RR*(*c*) = 1/*rank*(*c*), where *rank*(*c*) is the position of the first correct target in 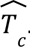. If no correct target appears within Top-k, *RR*(*c*) = 0. To match the cutoff used in the Magneto evaluation (Magneto-bp variants use the default topk = 20), we fix *K* = 20 for all *SchemaMapper* runs subjected to direct comparison against Magneto outputs.
- *Top-k accuracy. Top_k_(c)* = 1 if any element *G_c_* appears in the first *k* predictions, and 0 otherwise. We report k = 1-5.
- *Per-query Recall@GT.* Following the per-query formulation in Liu et al.^1^, 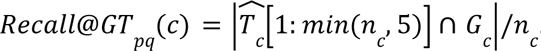. The truncation *min*(*n*_*c*ć_ 5) follows the released code and prevents the metric from rewarding methods solely for returning longer candidate lists.

#### Global Recall@GT

Whereas Top-k and MRR depend only on the ordering of candidates within each query and therefore have no analogous “global” form, Recall@GT can be computed against a study-wide ranked list of (query, target) pairs scored across all queries. The published Magneto results report a *global* Recall@GT, computed by flattening each study’s full source–target similarity matrix (up to 165 × 736 for this benchmark) into a list of (*c, t, s_c,t_*) triples ranked by score *s_c,t_*, retaining the top 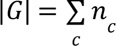 triples, and reporting the fraction that are ground-truth pairs. Unlike per-query Recall@GT, the global formulation requires that every (*c*, *t*) pair receive a comparable score, thereby penalizing cross-query score miscalibration: a method that ranks correct matches highly within each query but inconsistently across queries can score well per-query yet poorly globally.

*SchemaMapper* produces a sparse output by design: it roughly scores 5–20 candidates per query rather than the full target schema. Global Recall@GT is therefore not computable for *SchemaMapper* without modifying the architecture to score every (*c*, *t*) pair. We restrict *SchemaMapper* to per-query Recall@GT and report both formulations side by side for the Magneto-bp variants where both are available, leaving cells blank where the underlying scoring data does not support a given metric (**Table 1**).

#### Macro vs. micro aggregation

The published Magneto results are reported as macro averages, so we adopt macro averaging throughout the cross-method comparison in Table 1 to match the source paper’s aggregation convention. With n = 165 source columns distributed unevenly across 10 studies (n = 5-29), the choice of aggregation matters. Micro and macro averages of Top-k for the same *SchemaMapper* + Haiku-4.5-alias run differed by 1.73 pp at Top-1 and 0.44 pp at Top-5; the corresponding gap at MRR was the difference between micro (0.771) and macro (0.763) at Top-5. Mixing aggregation schemes across methods would introduce a systematic bias of comparable magnitude to the smallest method differences we report. The applicability of each aggregation scheme to each metric is summarized in **Supplementary Table 14**.

#### Cross-method comparability

MRR is computed identically for all four Magneto variants and both *SchemaMapper* configurations (Top-20 ranked list, macro-averaged across the 10 studies), and is therefore directly comparable across all six rows in Table 1. Top-1 and Top-5 accuracy are computed identically for *SchemaMapper* and the two Magneto-bp variants we reproduced locally; the Magneto-llm rows are not reported at the rank-distribution level needed to recompute these values without rerunning the proprietary LLM reranker. Per-query Recall@GT is comparable across the four reproducible rows (*SchemaMapper* variants and Magneto-bp variants), and global Recall@GT is comparable across the four Magneto rows (paper-reported and locally reproduced). We populate each cell only when the underlying scoring data support the metric and the formulation is consistent with the comparison group, and leave cells blank rather than substituting numerically incomparable values. Reproduction fidelity for the Magneto-bp variants is reported in the Table 1 caption.

#### Cascade-dependent rank behavior

The *SchemaMapper* cascade decides when to advance between stages based on the Top-1 confidence and merges results from each stage using Top-k-dependent rules. As a result, requesting more candidates does not simply extend the same ranked list – it can change which predictions appear in the Top-k, and the Top-5 of a k = 20 run is not in general identical to the Top-5 of a k = 5 run. To eliminate this confound from the cross-method comparison, all *SchemaMapper* numbers reported against Magneto are computed from a single k = 20 run, matching Magneto’s topk = 20 setting.

#### Confidence calibration

Confidence calibration metrics (AUC, Cohen’s *d*, Wilcoxon rank-sum) are computed directly over a pool of (prediction, correctness, confidence) records aggregated across all studies, rather than by aggregating per-query values. Because they have no per-query form, the macro/micro distinction does not apply (**Supplementary Table 14**). Within-study computation is degenerate for the smallest studies (e.g., Krug, *n* = 5), where the positive or negative class may be empty, and AUC is undefined; even when both classes are populated, sample sizes below ∼15 yield estimates with prohibitively wide confidence intervals. Pooling across studies avoids both failure modes and is the formulation we report.

### Denominator conventions across analyses

The four *OntologyMapper* benchmark inputs are not uniform in row count: UKBB-EFO and Biomappings-EFO have one row per query (888 and 795, respectively), while OLS-EFO (disease) and OLS-EFO (full) contain multi-target rows – the same source string appears multiple times with different ground-truth EFO labels – yielding 5,824 and 7,504 input rows, only 5,770 and 7,377 unique query strings, and 5,804 and 7,445 output rows. We adopt three different denominators across the analyses below, each matching the unit the corresponding metric is intended to summarize.

***Runtime analysis*** uses the **input benchmark row count** (888 / 795 / 5,824 / 7,504) because that is the workload each tool was handed. This measure answers “how long does the workflow take”, instead of “how fast is the matching algorithm itself”, which can be answered by using unique-query as the denominator. Both *OntologyMapper* and *text2term* internally deduplicate queries before invoking the matching backend, but the user-observable runtime (and the resulting output row count, after both tools expand results back to benchmark scale) is keyed to the input. The input row count, therefore, reflects the work actually executed and yields directly comparable per-query throughput across tools.

***Confidence-calibration analysis*** (mean confidence by correctness, Cohen’s d, AUC for correct-vs-incorrect separation) uses the **module output row count** (890 / 795 / 5,804 / 7,445 for *OntologyMapper*). Each emitted row is an independent prediction with its own Top-1 score; the calibration question is row-level (“do correct predictions score higher than incorrect ones?”), so collapsing rows would discard signals. *OntologyMapper* occasionally emits byte-identical duplicate rows for the same query (2 in UKBB; 34 in OLS-disease; 68 in OLS-full); these are retained because they represent independent module evaluations.

***Top-K accuracy and MRR*** use the **unique-query count** (888 / 795 / 5,770 / 7,377 for *OntologyMapper*; 888 / 795 / 5,824 / 7,504 for *text2term*). Accuracy answers a query-level question, “among the distinct mappings we wanted, what fraction landed in the Top-K?”. *OntologyMapper* deduplicates queries before Stage 2, so multi-target benchmark rows that don’t get resolved at Stage 1 collapse to a single matcher evaluation; *text2term* deduplicates before its TF-IDF call and then left-joins results back onto the full benchmark, so its denominator matches the raw benchmark row count. We report these conventions each tool naturally produces rather than forcing a common denominator, since either choice introduces an artificial bias. Query-count differences between tools (0–127 rows per benchmark) therefore reflect divergent handling of benchmark rows that share a query string but map to different targets, rather than differences in evaluation rigor. For all three analyses, ranking is computed case-insensitively, matching *text2term*’s built-in evaluation logic.

#### Statistical analysis for LLM-generated alias comparison

All comparisons are paired on the same 165 queries. For Top-k accuracy, we used McNemar’s exact two-sided test on the contingency of (rank ≤ k) indicators between method pairs. For MRR and Recall@GT, we used the paired Wilcoxon signed-rank test on per-query metric values. For each metric, we performed 28 pairwise comparisons across the eight methods. p-values were corrected for multiple comparisons by Holm–Bonferroni within each metric (m = 28). Where a Tier-A subset analysis is reported, Holm–Bonferroni was applied to the 15 within-Tier-A pairs (m = 15) as a planned-subset correction. Paired bootstrap confidence intervals on per-method MRR/Top-1/Top-5 and on the pairwise ΔMRR / ΔTop-k vs. Opus 4.5 were computed from B = 10,000 resamples (set.seed(42)); each bootstrap resample used the same query indices for every method to preserve pairing. A method’s CI on *‘Δ vs. Opus 4.5’* was classified as “statistically equivalent to Opus 4.5” when it crossed zero, and as “significantly worse” otherwise (no method had a CI lying entirely right of zero).

#### LLM-only baseline for schema mapping

The prompting configuration was driven by a common task-specific system instruction for schema matching (*”You are a biomedical data expert. Your task is to match a source metadata column to the most appropriate target column from a standardized schema. Return ONLY a JSON array; no prose, no markdown.”*) and a shared user template that listed all target columns and requested the Top-5 matches as a JSON array of {target, confidence} objects. The returned target was checked for membership in the target schema, and predictions outside the schema were labeled as hallucinations. Top-k accuracy (k = 1,3,5), MRR, and Recall@GT were computed; 95% confidence intervals on Top-k accuracy were bootstrap replicates (n = 1,000). Paired comparisons between configurations use the Wilcoxon signed-rank test on per-query rank differences (one-sided). For Top-1 correctness, we additionally report discordant-pair (McNemar-style) counts with an exact binomial test against p = 0.5. Five frontier models spanning two independent pretraining corpora were evaluated on GDC: Claude Haiku 4.5, Claude Sonnet 4.5, and Claude Opus 4.5 (Anthropic), as well as Gemini 2.5 Pro and Gemini 2.5 Flash (Google). All calls used temperature = 0, where the provider permits. Opus 4.7 was used for *SchemaMapper* alias generation but was excluded from LLM-only because it does not accept temperature = 0. Requests were issued sequentially at 1s intervals with exponential backoff retries. All responses were cached on disk to make re-runs deterministic.

#### LLM-only baseline for ontology mapping

An LLM-only baseline was evaluated on the 888-query UKBB-EFO benchmark using the EFO corpus (17,638 terms from EFO v3.62.0). Unlike the corpus used for benchmarking against *text2term*, it includes only EFO-native classes, and imported, non-EFO classes (e.g., from HP, MONDO, etc.) are excluded. We used this smaller corpus because the 33,230-term EFO corpus exceeds Haiku 4.5’s allowed input token size. Two prompting configurations were compared: 1) *zero_shot* - a role instruction only with the user message containing only the query phrase and a request for the Top-5 EFO labels; 2) *open_book_full* - the full EFO corpus injected as a cached system block with Anthropic’s ephemeral prompt caching, with a “verbatim from the list” constraint in the user message. The ∼161k-token corpus block is billed once upon first creation and then read at ∼10% of the base input rate for the remainder of the ephemeral TTL (Time to Live), making the full 888-query pass affordable. Each returned label was checked against the 16,218 unique EFO labels (17,638 terms without 1,419 obsolete terms and 1 duplicated term). Predictions absent from it were labeled as hallucinations. Accuracy at Top-k and bootstrap 95% CIs were computed in the same way as in the main benchmarking. Claude Haiku 4.5 was used with temperature = 0, max_tokens = 1024, sequential requests at 1s intervals, and up to 5 exponential-backoff retries for transient errors. All responses were cached on disk.

#### LLM contamination control for schema mapping

Because the GDC data model and CPTAC harmonization mappings are openly documented and present in common web-scale pretraining corpora, strong LLM-only performance on the GDC benchmark may reflect memorization of target-schema identifiers rather than transferable schema-matching capability. To quantify the contribution of memorization, we applied a three-part battery (E2, E3, E4) to each configuration on the GDC benchmark in addition to the baseline (E1). We ran identical perturbations through three configurations: 1) *SchemaMapper* without an alias dictionary, a pure embedding-plus-retrieval pipeline referred to below as ‘*SM-no-alias*’, 2) *SchemaMapper* with each of six LLM-generated alias dictionaries (*SM+alias*; Haiku 4.5, Sonnet 4.5, Opus 4.5, Opus 4.7, Gemini 2.5 Flash, Gemini 2.5 Pro), and 3) *LLM-only*. For LLM-only, we evaluated the same five frontier models defined in the LLM-only baseline for schema mapping (**Supplementary Methods**): Opus 4.7 was excluded because it rejects temperature=0 and would not have produced comparable deterministic outputs.

***Source-side perturbations (E2)*** rewrite the source column names without changing ground truth. Synonym-swap uses Claude Sonnet 4.5 (temperature = 0) to generate a snake_case paraphrase, rejecting and regenerating any output that matches a GDC target name exactly (up to two retries). Abbreviation-flip expands abbreviated source names to full form and vice versa. A 20% random sample (25 of 124 unique sources) was audited for semantic preservation. Crucially, for the *SM+alias* configurations under E2, the original alias dictionaries were retained unchanged.

### The target-side perturbation (E3)

All 736 GDC target column identifiers were renamed to semantically equivalent non-GDC-style strings using Sonnet 4.5 (0 empty outputs; 0 collisions with original GDC names; 147 / 736 rows audited); thus, the semantic matching capability is preserved by construction. Ground truth was translated through the rename map, so a “correct” prediction under E3 is the renamed label for the same source concept. For the *SM+alias* configurations, the alias dictionary’s field_name column was renamed, so a ‘correct’ answer under E3 is the renamed target. Any perturbation output that exactly matches a GDC target name was rejected (case-insensitive, up to 2 retries) to prevent LLM-only from using memorized mapping. The E3 paired comparison is computed on the subset of unique source columns (n = 124) rather than the full 165-row benchmark, because several source columns recur across CPTAC studies and would otherwise contribute correlated pairs that violate the independence assumption of the paired Wilcoxon test. E1 baseline accuracies are recomputed on the same unique subset, so the comparison is apples-to-apples. Qualitative conclusions are unchanged.

#### Direct memorization probes (E4)

Independent of the schema-matching task, we employed three diagnostic tests (probes, *P*) to test for memorized GDC structure. *P1* asks each model to list, from zero context, all columns of the GDC demographic node and scores the fraction of output lines whose first token is a real GDC column. *P2* asks for the exact GDC identifier for each of 20 common clinical concepts (“biological sex”, “AJCC pathologic tumor stage”), with no schema provided, and reports the exact-match rate. *P3* presents a 12-name slice of the GDC schema, with three names randomly masked as ???, and asks the model to fill them in (10 slices, seeded); we report the exact recovery rate.

#### Duplicate-query handling

The 165-mapping benchmark contains 124 unique source column names; 26 source strings recur across multiple CPTAC studies (e.g., ‘Stage’ appears in three studies, ‘Tumor_Site’ in two). These are not redundant evaluations: the gold-standard ‘ref_match’ is defined per source study, so the same column name can resolve to different harmonized GDC targets across studies. All Top-1 and MRR accuracies are reported at the row level over the full n=165 evaluations, preserving per-study labels. Paired comparisons between baseline (E1) and perturbed runs (E2, E3) are matched on the ‘(query, source_study)’ key rather than ‘query’ alone. LLM predictions are deterministic under ‘temperature=0’ and we confirmed that every duplicated source string produced identical Top-1 predictions across its copies in all E1/E2/E3 runs; nonetheless, correctness diverges across copies when the per-study gold differs.

## Supplementary Results

### *SchemaMapper* Alias Dictionary Generation Feasibility and Cost

Of the nine models tested, seven successfully completed the 736-field alias dictionary generation, achieving 100% field coverage; per-model row counts, wall times, and API costs are reported in **Supplementary Table 9**. Gemma3:27b required post-hoc filtering of 185 rows associated with 105 hallucinated field names (1.9% of generated output) before downstream use. Two open-weight models failed outright: Gemma4:26b produced malformed output that could not be parsed, while Qwen3:32b returned *structurally* valid output in which the source-alias column contained the literal letters A–F (the placeholder labels from the few-shot example table) rather than real alias strings, a failure mode sometimes referred to as template capture. Because the prompt was held fixed across all generators for parity, we did not attempt prompt-level rehabilitation of Qwen3:32b.

### Top-tier LLM-generated alias models are mutually interchangeable under family-wise correction; the equivalent set narrows under per-pair confidence intervals

Six of the seven alias models — Opus 4.5, Haiku 4.5, Gemini 2.5 Pro, Opus 4.7, Gemini 2.5 Flash, and Sonnet 4.5 (henceforth ‘Tier-A’) — significantly outperformed the no-alias *SchemaMapper* baseline on Top-1 after Holm–Bonferroni correction over the 28 all-pairs McNemar comparisons (Holm-adjusted p ≤ 0.032). Gemma 3 27B did not (Holm-p = 1.0; raw p = 0.54).

#### Family-wise comparison, answering “is any Tier-A pair reliably different from any other?”

Within Tier-A, all 15 within-tier comparisons were mutually non-significant under the same 28-pair family (Holm-adjusted McNemar p ≥ 0.105 within the Tier-A 15 pair family, p ≥ 0.119 across all 28 pairs). The same conclusion held for Recall@GT (Holm-adjusted paired Wilcoxon signed-rank p ≥ 0.273 within Tier-A, p ≥ 0.291 across all 28 pairs). MRR was the most sensitive continuous metric, identifying two within-Tier-A pairs as significant (Opus 4.5 > Sonnet 4.5, Holm-p = 0.004; Opus 4.5 > Gemini 2.5 Flash, Holm-p = 0.042) and two as borderline (Opus 4.5 > Opus 4.7 and Haiku 4.5 > Sonnet 4.5, Holm-p = 0.052 each). Top-3 and Top-5 each identified one within-Tier-A pair as significant, Opus 4.5 > Opus 4.7 (Top-3 Holm-p = 0.026; Top-5 Holm-p = 0.003). Notably, Opus 4.5 vs. Haiku 4.5 was statistically indistinguishable on every metric (Holm-p = 1.0 across Top-1, Top-3, Top-5, MRR, and Recall@GT).

#### Per-pair comparison, answering “can any single model be comparable to Opus 4.5 as a deployment option?”

Per-pair 95% paired-bootstrap CIs on Δ vs. Opus 4.5 are stricter than the Holm-corrected pairwise test, and the equivalent set narrows as the metric becomes more discriminative (**Supplementary Figure 1A-C**). The ΔMRR CI crosses zero only for Haiku 4.5 and (Δ = −0.029, CI [−0.078, +0.019]) and Gemini 2.5 Pro (Δ = −0.041, CI [−0.091, +0.006]). The ΔTop-1 CI additionally includes Opus 4.7 (Δ = −5.5 pp, CI [−11.5, +0.6 pp]), though this is the most fragile equivalence call, with the upper bound only just crossing zero. At Top-5, the equivalent set collapses to just Opus 4.5 and Haiku 4.5. The ΔRecall@GT CI crosses zero only for Haiku 4.5 and Gemini 2.5 Pro.

### Two Tier-A models are strictly Pareto-dominated

Plotting Top-1 / MRR against one-time generation cost revealed the Pareto frontier of cost-efficient alias models (**Supplementary Figure 1D**). Among the API-hosted methods, the frontier (in ascending cost) consists of Gemini 2.5 Flash ($0.49) → Haiku 4.5 ($1.04) → Gemini 2.5 Pro ($2.00) → Opus 4.5 ($21.51). Two Tier-A methods are off-frontier and strictly dominated by the same-vendor alternative: Sonnet 4.5 ($4.45, MRR 0.708) is dominated by Haiku 4.5 ($1.04, MRR 0.771): the same vendor, ≈ 4× more expensive, and lower MRR. Opus 4.7 ($28.13, MRR 0.730, 2.5h generation) is dominated by Opus 4.5 ($21.51, MRR 0.800, 0.8h generation) (**Supplementary Table 9**): the same vendor, more expensive, slower, and lower MRR on every metric, with the Top-3/Top-5 gap reaching statistical significance.

### Generalizability limitations

Several caveats constrain the generalizability of these findings. First, the benchmark scope was limited; n = 165 paired queries spread across ten CPTAC studies provide limited statistical resolution. Second, the benchmark was performed over a single domain. The CPTAC studies are exclusively cancer proteomics, and the alias-generation prompt was the cancer_genomics prompt tuned for that domain. Whether the same Tier-A equivalence holds for non-oncology biomedical metadata (e.g., microbiome, EHR, ecology) remains unverified. Cross-domain replication is essential before generalizing the “pick on cost” recommendation outside oncology genomics. Lastly, our statistical tests assumed per-query independence. McNemar’s test, the Wilcoxon signed-rank test, and the paired bootstrap all assume independent paired observations. The 66 multi-GT queries (G > 1) violate this mildly — they contribute correlated within-query observations to Recall@GT. We accept this as a small bias against the conservative test direction and do not attempt a hierarchical correction.

### Source-side paraphrase reveals a robustness gap in *SchemaMapper*, not an LLM strength

The source-side perturbation (E2) showed the opposite asymmetry to the target-side rename (E3) and identified the one aspect where LLM is more robust than the *SchemaMapper* architecture. Under synonym-swap of source column names, LLM-only lost 7.9–11.5 pp Top-1, whereas every SM+alias configuration lost substantially more (22.4–29.7 pp), and SM-no-alias fell from 53.9 → 33.3% (−20.6 pp, **Supplementary Table 12**): LLMs paraphrase-match degrade more gracefully, while *SchemaMapper*’s embedding retrieval is more sensitive to the lexical form of the source column. Abbreviation-flip followed the same pattern but with smaller magnitudes (LLM-only losses 0.4–6.5 pp; *SchemaMapper* losses 9.1–16.4 pp), consistent with medical abbreviations being sufficiently standardized that both surface forms are equally familiar to both model families and both embedding-based retrievers. Critically, for the SM+alias configurations under E2, the original alias dictionaries were retained unchanged, so these losses reflect the matching stage’s sensitivity to source paraphrase rather than degraded alias coverage. This is a limitation of the current *SchemaMapper* retrieval rather than evidence of LLM task competence: unlike the E3 target-rename result, the E2 gap does not invert, and it points to source-side normalization or paraphrase-aware query expansion as a concrete direction for improving *SchemaMapper* robustness.

## Supplementary Figures

**Supplementary Figure 1.**
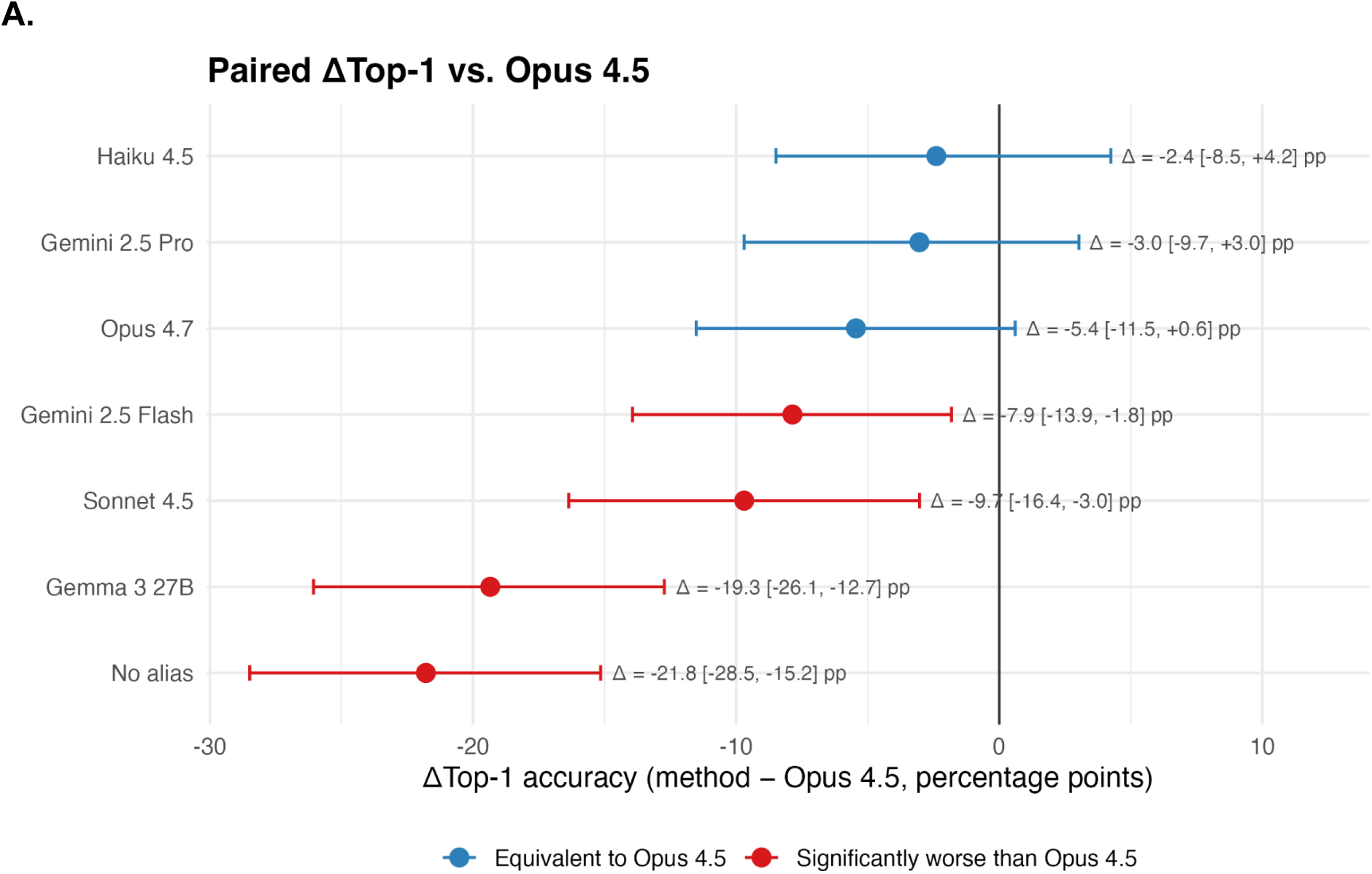
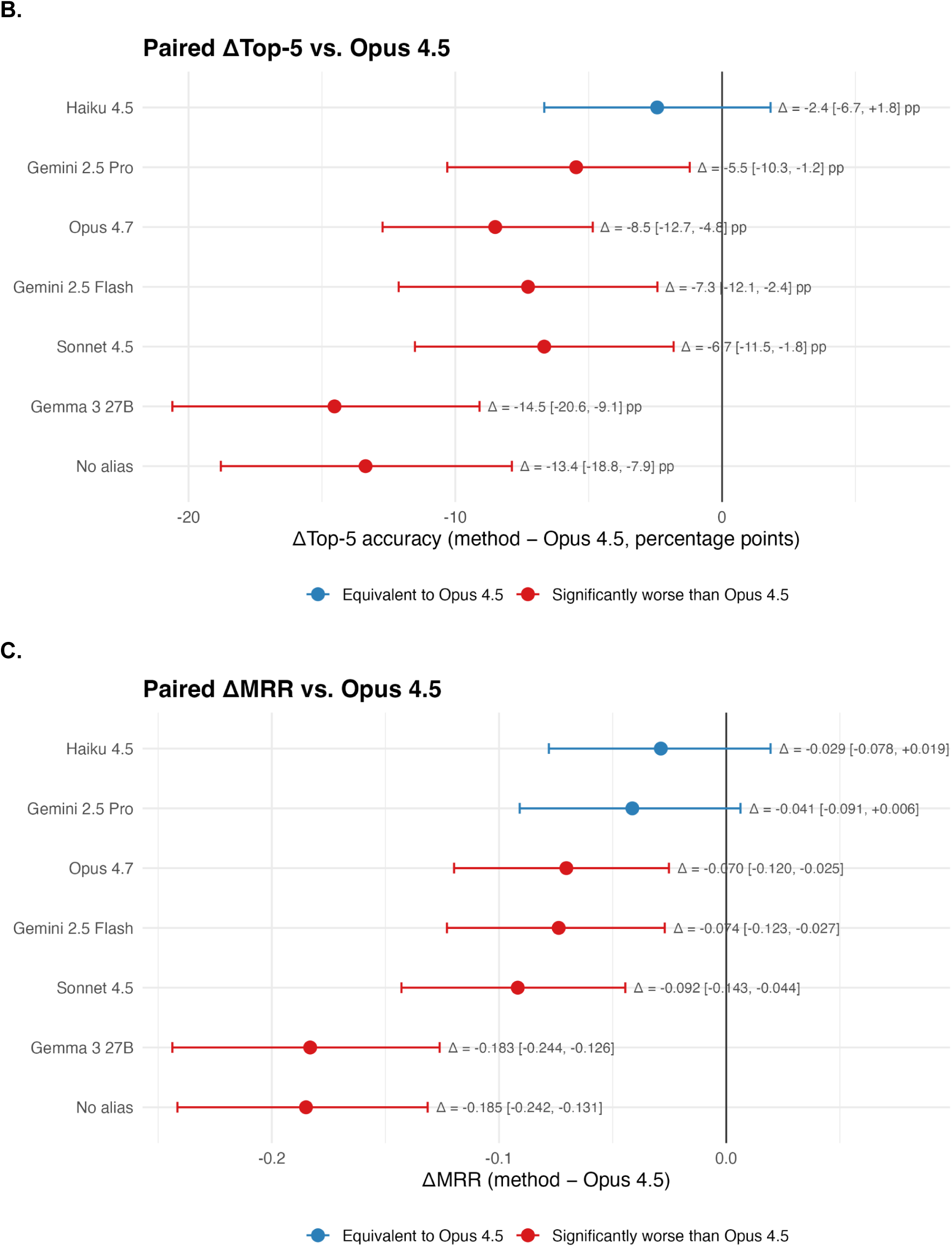
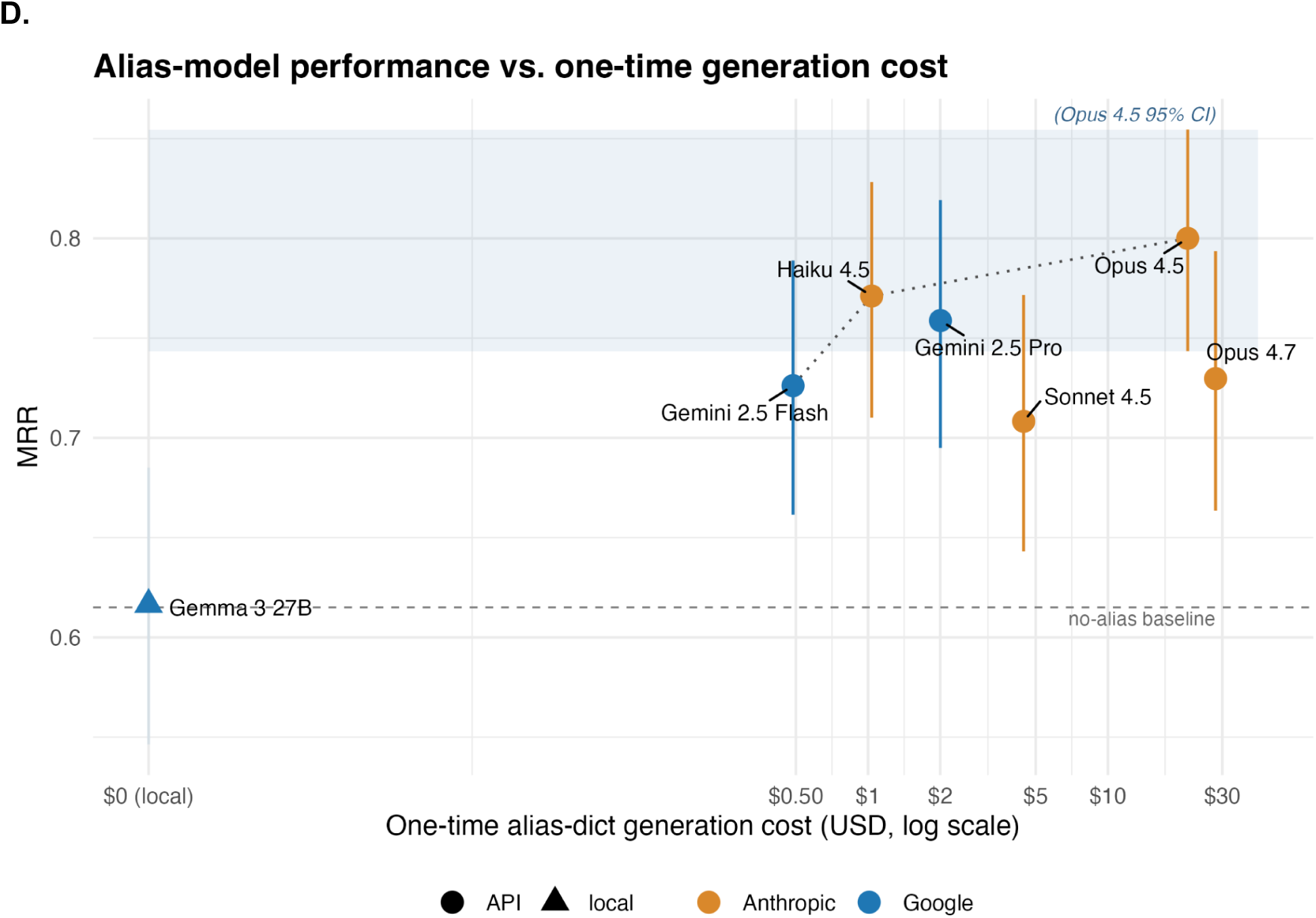
Choice of LLM model for *SchemaMapper* alias generation. Each alias dictionary was generated by a single LLM and supplied to Stage 1 of *SchemaMapper*, and performance was evaluated on the GDC schema-mapping dataset as in Figure 2. Panels A–C show paired-bootstrap comparisons of each dictionary against the Opus 4.5 dictionary (Δ = method − Opus 4.5), with 95% CIs from B = 10,000 query-matched resamples (n = 165). *Methods whose CI crosses zero (blue) are statistically equivalent to Opus 4.5*; methods whose CI lies entirely below zero (red) are significantly worse. **(A)** ΔTop-1 accuracy. Haiku 4.5, Gemini 2.5 Pro, and Opus 4.7 cross zero, though the Opus 4.7 interval only marginally includes zero. **(B)** ΔTop-5 accuracy. Only Haiku 4.5 was equivalent to Opus 4.5, while all other dictionaries were well below zero. **(C)** ΔMRR. Haiku 4.5 and Gemini 2.5 Pro cross zero. Across panels, the equivalent-to-Opus-4.5 set narrows as the metric becomes more rank-sensitive (Top-1 → MRR → Top-5), with Haiku 4.5 the only dictionary indistinguishable from Opus 4.5 on every metric. **(D)** MRR (95% paired-bootstrap CI) against one-time alias-dictionary generation cost (USD, log scale). The shaded band is the Opus 4.5 95% CI, and the dashed line is the no-alias baseline. Point shape denotes hosting (circle = API, triangle = local) and color denotes vendor (orange = Anthropic, blue = Google). The dotted line traces the cost–performance Pareto frontier (Gemini 2.5 Flash → Haiku 4.5 → Gemini 2.5 Pro → Opus 4.5); Sonnet 4.5 and Opus 4.7 sit off-frontier, each dominated by a cheaper same-vendor alternative.

## Supplementary Tables

**Supplementary Table 1.**
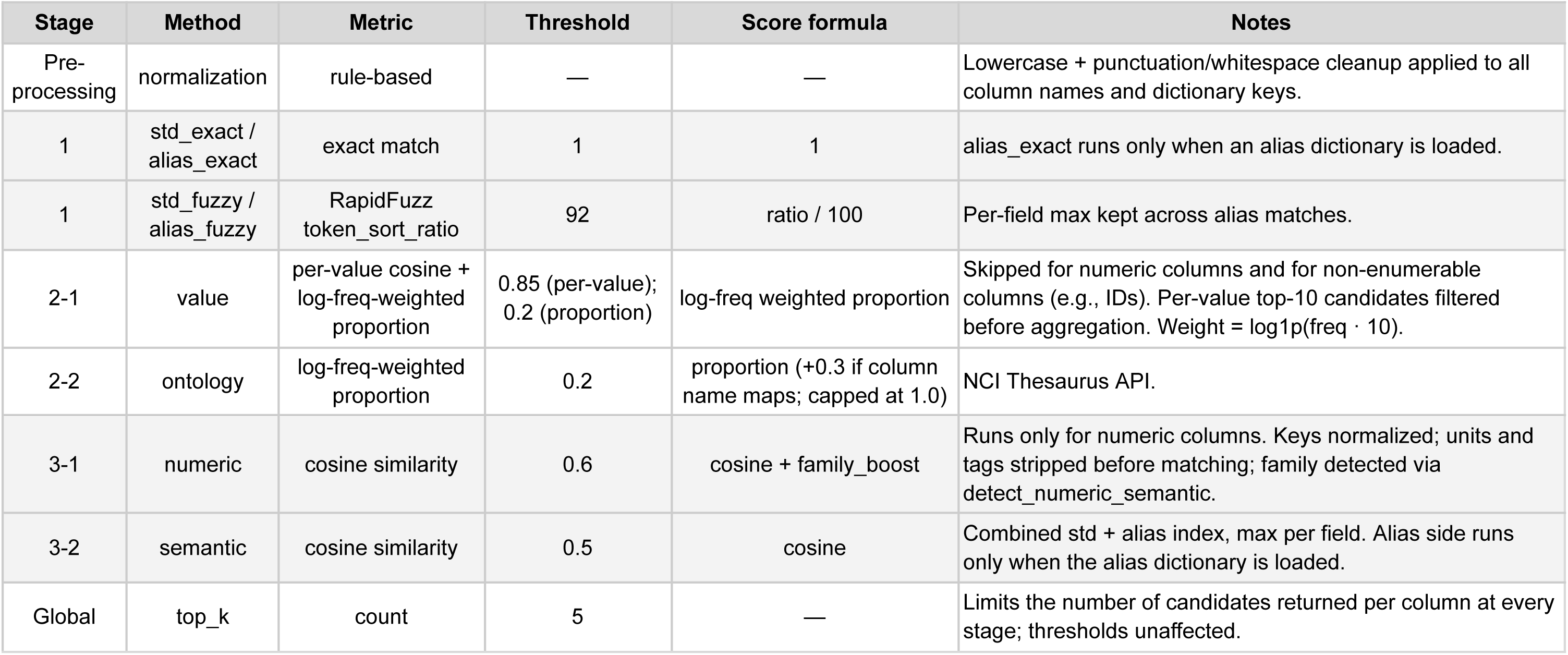
*SchemaMapper* configuration parameters. Per-stage thresholds, methods, and acceptance rules. All embedding-based methods use the all-MiniLM-L6-v2 sentence encoder by default; Stage 2 ontology matching additionally uses the NCI Thesaurus API. Cascade behavior across stages: each method accepts when its Top-1 score meets the threshold; otherwise, the best-seen candidate is retained and the cascade advances. Sub-threshold results are still returned if no better match is found.

**Supplementary Table 2.** *OntologyMapper* configuration parameters. Per-stage thresholds, methods, and acceptance rules. Embeddings use SapBERT (cambridgeltl/SapBERT-from-PubmedBERT-fulltext) by default. All FAISS indices are IndexFlatIP over L2-normalized vectors, making the inner product equivalent to cosine similarity. Cascade behavior: Stage 1 exact matches bypass all later stages; non-exact queries enter Stage 2 unconditionally; Stage 2.5 is invoked only when the Stage 2 Top-1 score falls below the synonym-boost threshold.

| Stage | Method | Metric | Threshold | Score formula | Notes |
| --- | --- | --- | --- | --- | --- |
| 1 | exact | case-insensitive string match | 1 | 1 | .strip().lower() against corpus set. |
| 2 | lm (default) | FAISS inner product | — | FAISS IP | CLS-token embedding via HuggingFace AutoModel; max_length = 128. |
| 2 | st | FAISS inner product (cosine-equivalent) | — | cosine | SentenceTransformer pooling per checkpoint; normalize_embeddings = True. |
| 2.5 | synonym boost | synonym lookup score | 0.9 | max(Stage 2 score, synonym score) per term | Targets Stage 2 rows with top-1 < 0.9. Merges Stage 2 and synonym candidates, max score per term, re-ranks Top-k. Stage label updated to 2.5 only if boosted (new Top-1 > old Top-1, or match_level improves). Synonym tables sourced from NCI EVSREST or OLS per ontology_source. |
| Global | top_k | count | 5 | — | Limits the number of returns per query at every stage. |
| Global | filter_obsolete | prefix filter | False (default) | — | When True, drops obsolete_* labels from both corpus and synonym candidates. |
| Global | corpus registry | lookup | — | — | Auto-resolves (category, ontology_source) to root term; e.g., (disease, ncit) → NCIT:C3262. Other combinations require a user-supplied corpus. |

**Supplementary Table 3.**
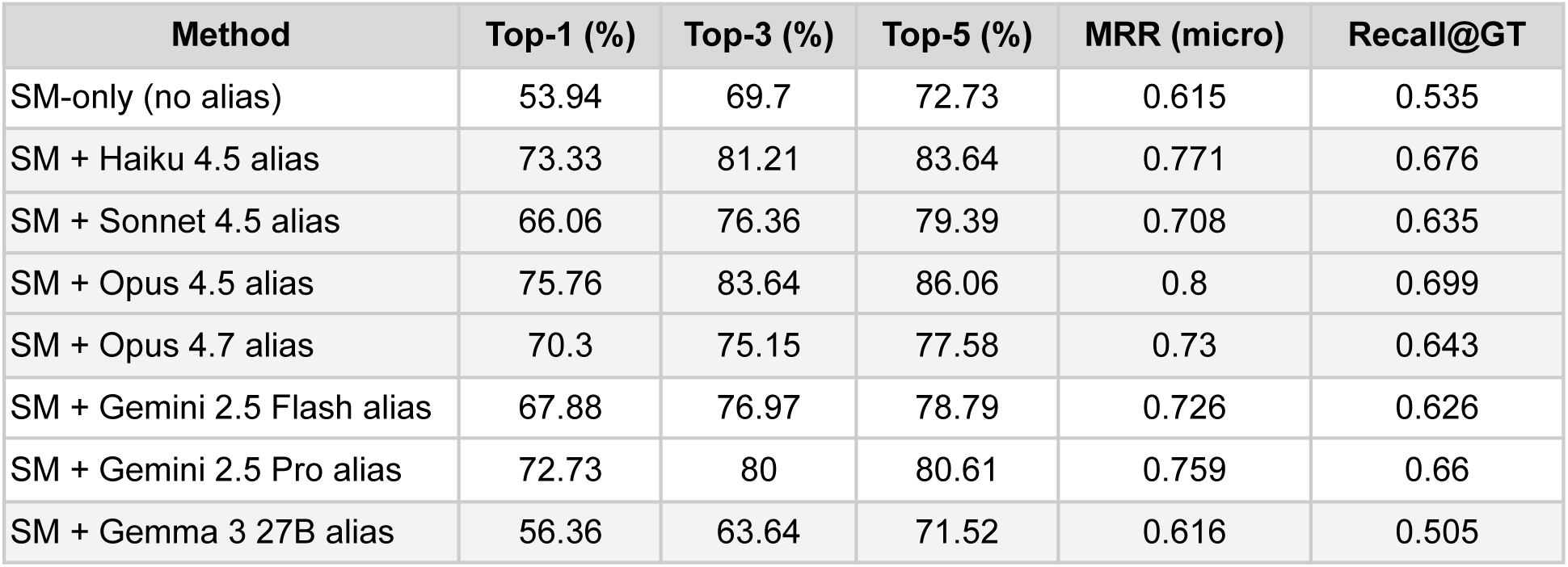
*SchemaMapper* performance under different alias-dictionary configurations. We compared the performance of five alias variants for schema mapping across 10 CPTAC studies. Variants differ only in the alias dictionary supplied to Stage 1 of the *SchemaMapper* pipeline: seven LLM-generated dictionaries and a no-alias control on the raw column names only. All metrics are per-query means over the 165 ground-truth mappings (case-insensitive matching). Micro-average is used here; by contrast, Table 1 uses macro-averaging to match the Magneto convention. SM = *SchemaMapper*.

**Supplementary Table 4.** *SchemaMapper* performance by pipeline stage and method, with or without LLM-generated aliases. Each row reports the number of queries (*n*) resolved at a given stage/method combination across all 10 CPTAC studies (165 queries total), along with the corresponding accuracy at rank 1, 3, and 5 (Top-k) and Mean Reciprocal Rank (MRR). Invalid queries (4 for Stage 1 and 13 for Stage 3) contribute zeros to all accuracy metrics. The “Haiku-4.5 alias” configuration adds a Haiku 4.5–generated alias dictionary on top of the target dictionary; “No alias” uses the target dictionary only. Adding aliases reroutes 30 queries from Stage 3 to Stage 1 (27 alias_exact + 3 alias_fuzzy) and improves the residual Stage 3 semantic retrieval (Top-1 38.6% → 62.1%; MRR 0.495 → 0.692), indicating that LLM aliases both deflect easy cases away from the semantic stage and leave a more tractable residual for embedding-based matching. We used the Top-5 retrievals and the micro-average for this evaluation.

| Configuration | Stage | Method | n | Top-1 (%) | Top-3 (%) | Top-5 (%) | MRR (micro) |
| --- | --- | --- | --- | --- | --- | --- | --- |
| Haiku-4.5 alias | Stage 1 | std_exact | 46 | 95.7 | 95.7 | 95.7 | 0.957 |
|  | Stage 1 | std_fuzzy | 1 | 100 | 100 | 100 | 1 |
|  | Stage 1 | alias_exact | 27 | 81.5 | 81.5 | 81.5 | 0.815 |
|  | Stage 1 | alias_fuzzy | 3 | 0 | 0 | 0 | 0 |
|  | Stage 2 | value/ontology | 1 | 0 | 0 | 0 | 0 |
|  | Stage 3 | semantic | 87 | 62.1 | 77 | 81.6 | 0.692 |
| No alias | Stage 1 | std_exact | 46 | 95.7 | 95.7 | 95.7 | 0.957 |
|  | Stage 1 | std_fuzzy | 1 | 100 | 100 | 100 | 1 |
|  | Stage 2 | value/ontology | 4 | 0 | 0 | 0 | 0 |
|  | Stage 3 | semantic | 114 | 38.6 | 61.4 | 65.8 | 0.495 |

**Supplementary Table 5.** Per-stage accuracy of *OntologyMapper* on the four EFO benchmarks. Each query is attributed to the pipeline stage that produced its Top-1 prediction. Per-stage *n* are output-row counts (see Denominator conventions, **Supplementary Methods**); these differ slightly from the unique-query denominators used in Table 2, so per-stage and overall accuracies rest on marginally different bases. Stage 1 is exact-match by construction, so its Top-K is 100%. We used the Top-5 retrievals and the micro-average for this evaluation.

| Benchmark | Stage | n | Top-1 (%) | Top-3 (%) | Top-5 (%) | MRR (micro) |
| --- | --- | --- | --- | --- | --- | --- |
| UKBB-EFO | 1 | 288 | 100 | 100 | 100 | 1 |
|  | 2 | 284 | 70.42 | 82.04 | 84.51 | 0.763 |
|  | 2.5 | 318 | 65.41 | 80.82 | 85.22 | 0.733 |
| Biomappings-EFO | 1 | 576 | 100 | 100 | 100 | 1 |
|  | 2 | 127 | 85.04 | 94.49 | 97.64 | 0.906 |
|  | 2.5 | 92 | 81.52 | 92.39 | 92.39 | 0.861 |
| OLS-EFO (disease) | 1 | 4,014 | 100 | 100 | 100 | 1 |
|  | 2 | 1,206 | 88.64 | 95.52 | 96.77 | 0.921 |
|  | 2.5 | 584 | 75.68 | 83.05 | 85.79 | 0.796 |
| OLS-EFO (full) | 1 | 4,825 | 100 | 100 | 100 | 1 |
|  | 2 | 1,552 | 76.1 | 82.02 | 83.18 | 0.791 |
|  | 2.5 | 1,068 | 59.18 | 67.23 | 69.57 | 0.634 |

**Supplementary Table 6.** Confidence calibration of *OntologyMapper* Top-1 similarity scores across four EFO benchmarks. For each benchmark, queries are partitioned into correctly and incorrectly mapped at Top-1 (n correct, n incorrect). Mean conf. (correct) and Mean conf. (incorrect) are the average Top-1 cosine similarities in each group; Δ is their difference (correct − incorrect). Cohen’s d quantifies the standardized separation between the two confidence distributions (pooled SD), and AUC is the area under the ROC curve for using the Top-1 score as a classifier of mapping correctness (0.5 = chance, 1.0 = perfect separation). Larger Δ, |d|, and AUC indicate that the Top-1 score is more informative as a confidence signal (i.e., better-calibrated). All four benchmarks use the baseline configuration (Stages 1/2/2.5 with SapBERT embeddings). Wilcoxon rank-sum p-values were < 2 × 10⁻¹⁶ for all rows (Holm-adjusted). We used the Top-5 retrievals and the micro-average for this evaluation.

| Benchmark | n correct | n incorrect | Mean conf. (correct) | Mean conf. (incorrect) | $\Delta$ | Cohen's d | AUC |
| --- | --- | --- | --- | --- | --- | --- | --- |
| UKBB-EFO | 694 | 196 | 0.943 | 0.86 | 0.0829 | 0.954 | 0.766 |
| OLS-EFO (full) | 6,638 | 807 | 0.988 | 0.851 | 0.1368 | 2.579 | 0.895 |
| OLS-EFO (disease) | 5,525 | 279 | 0.99 | 0.83 | 0.1602 | 3.89 | 0.935 |
| Biomappings-EFO | 759 | 36 | 0.986 | 0.871 | 0.1148 | 2.804 | 0.917 |

**Supplementary Table 7.** *OntologyMapper* runtime comparison between cold (first-time) and warm (cached) runs. We measured *OntologyMapper’s* runtime on the 33,230-term EFO corpus. Cold and warm runs are defined in the Methods section. ***Init***: concept table construction time. ***Model Init***: time to load SapBERT model weights from the local cache directory into memory. ***Inference***: Stage 2 SapBERT forward pass, comprising query embedding and FAISS nearest-neighbor search against the corpus. ***Pipeline***: cumulative wall-clock time across all mapping stages (normalization, Stages 1, 2, and 2.5), excluding initialization. ***Total***: Init + Pipeline, representing the full end-to-end runtime from invocation to completion. All times are reported in seconds.

| Run | Benchmark | n | Init (s) | Model Init (s) | Inference (s) | Pipeline (s) | Total (s) |
| --- | --- | --- | --- | --- | --- | --- | --- |
| Cold (first) | UKBB-EFO | 888 | **1168.3** | NA | NA | 11.7 | 1180 |
| Cold (first) | Biomappings-EFO | 795 | **1342.9** | NA | NA | 4.1 | 1347 |
| Cold (first) | OLS-EFO (disease) | 5824 | **1290.4** | NA | NA | 28.6 | 1319 |
| Cold (first) | OLS-EFO (full) | 7504 | **1279.1** | NA | NA | 42.9 | 1322 |
| Warm (cached) | UKBB-EFO | 888 | **0.36** | 0 | 9.71 | 11.75 | 12.11 |
| Warm (cached) | Biomappings-EFO | 795 | **0.33** | 0 | 3.37 | 4.12 | 4.45 |
| Warm (cached) | OLS-EFO (disease) | 5824 | **0.31** | 0.01 | 25.31 | 28.57 | 28.89 |
| Warm (cached) | OLS-EFO (full) | 7504 | **0.3** | 0 | 37.18 | 42.9 | 43.2 |

**Supplementary Table 8.** Wall-clock runtime of *OntologyMapper* and *text2term* across four EFO benchmarks. Per-benchmark execution time (mm:ss) for *OntologyMapper* across two operating modes, compared with *text2term* (TF-IDF baseline). *Cold* denotes the first run against a fresh corpus, including one-time construction of the embedding index over a 12-ontology EFO corpus (33,230 terms for *OntologyMapper* and 33,659 terms for *text2term*) parsed from a local OWL file and FAISS index serialization to disk. *Warm* denotes subsequent runs that reuse the cached FAISS index and SQLite metadata store, reflecting the steady-state cost users incur after initial setup. *text2term* (TF-IDF) timings include the local OWL parse; the original *text2term* evaluation pulled the OWL over HTTP, adding ∼5s per benchmark. *n* queries indicate the number of input strings mapped per benchmark. Totals are summed across all four benchmarks (15,011 queries). All measurements were collected on a single workstation: Apple M4 Pro, 48 GB unified memory, internal SSD, macOS Sequoia 15.6.1 (aarch64), with no concurrent workloads. Cold-mode cost is incurred once per (model, corpus-content) pair and is excluded from the typical end-user runtime once the index is cached.

| Benchmark | n queries | Cold | Warm | text2term |
| --- | --- | --- | --- | --- |
| UKBB-EFO | 888 | 19m 40s | 12.1s | 14.4s |
| Biomappings-EFO | 795 | 22m 27s | 4.5s | 13.8s |
| OLS-EFO (disease) | 5824 | 21m 59s | 28.9s | 14.8s |
| OLS-EFO (full) | 7504 | 22m 02s | 43.2s | 15.3s |
| **Total** | **15,011** | **1h 26m 08s** | **1m 28.7s** | **58.3s** |

**Supplementary Table 9.** Summary of LLM-generated alias dictionaries for the 736-field GDC schema. Nine large language models were evaluated for their ability to generate alias dictionaries for the 736 GDC-standard fields used in the *SchemaMapper* benchmark. Four Anthropic models (Claude Haiku 4.5, Sonnet 4.5, Opus 4.5, Opus 4.7), two Google API models (Gemini 2.5 Flash, Gemini 2..5 Pro), and three locally hosted open-weight models (Gemma3:27b, Gemma4:26b, Qwen3:32b) were run with an identical five-pass prompt template; no prompt-level adjustments were made per model. Rows is the total number of (field, alias) entries in the resulting dictionary after format validation and, for Gemma3:27b, post-hoc removal of hallucinated field names. Aliases per field are computed as Rows / 736, except for Claude Haiku 4.5, where the reported value reflects the per-field mean after de-duplication. Wall time is end-to-end generation time on the production hardware (Anthropic API for Claude models; local GPU for open-weight models). API cost is the total amount billed by the provider; locally run models incur no API charge. Failed runs (Gemma4:26b: malformed output that could not be parsed; Qwen3:32b: structurally valid output in which the source-alias column contained the literal placeholder labels A–F from the few-shot example, a “template capture” failure mode) produced no usable dictionary and are reported with em-dashes. All completed dictionaries achieved 100% field coverage.

| Model | Provider / Mode | Status | # of Alias | Aliases per field | Top-1 (%) | Wall time | API cost (USD) | Notes |
| --- | --- | --- | --- | --- | --- | --- | --- | --- |
| Claude Haiku 4.5 | Anthropic API | Completed | 9,219 | 12.4 | 73.33 | 16.5 min | ~\$1.04 | |
| Claude Sonnet 4.5 | Anthropic API | Completed | 14,048 | 19.1 | 66.06 | 49 min | \$4.45 | |
| Claude Opus 4.5 | Anthropic API | Completed | 14,200 | 19.3 | 75.76 | 46 min | ~\$21.51 | |
| Claude Opus 4.7 | Anthropic API | Completed | 12,515 | 17 | 70.3 | 2 h 31 min | \$28.13 | |
| Gemini 2.5 Flash | Google API | Completed | 9,440 | 13 | 67.88 | 13.4 min | ~\$0.49 | 151 rows from 7 hallucinated fields filtered (1.3% of raw) |
| Gemini 2.5 Pro | Google API | Completed | 10,756 | 15 | 72.73 | 41 min | ~\$2.00 | 2 rows from 2 hallucinated fields filtered (0.02% of raw) |
| Gemma3:27b | Open-weight (local) | Completed | 7,757 | 10.5 | 56.36 | 4 h 41 min | \$0.00 | 185 rows from 105 hallucinated fields filtered (1.9% of raw) |
| Gemma4:26b | Open-weight (local) | Failed | — | — | — | — | — | Failed: format compliance issue |
| Qwen3:32b | Open-weight (local) | Failed | — | — | — | — | — | Failed: template capture (literal A–F from few-shot example) |

**Supplementary Table 10.**
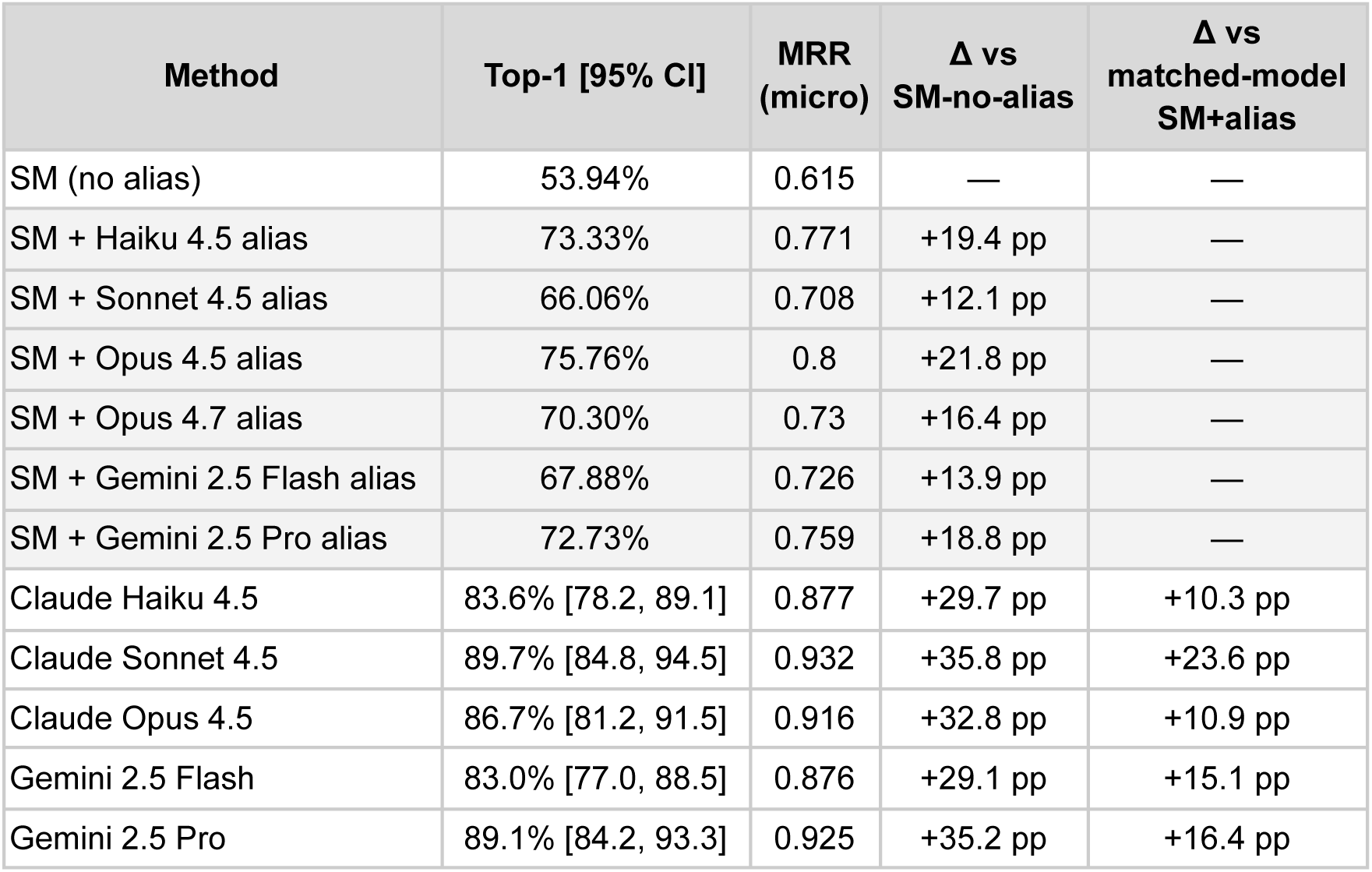
LLM-only baseline vs *SchemaMapper* on the GDC schema-mapping benchmark. We compared the schema-mapping performance of different configuration families on 165 source-to-target column mappings drawn from 10 CPTAC proteomics studies against the 736-column GDC clinical data model. Only the source and target column names were used. Three configuration families are: (i) *SchemaMapper* without alias enrichment (Stages 1–3); (ii) *SchemaMapper* with LLM-generated alias dictionaries (Stages 1–3 plus a Stage-1 alias dictionary of 9,000–14,000 entries, one dictionary per LLM family); and (iii) LLM-only, in which the source column is sent to a frontier LLM together with the full 736-column target list in a single zero-shot call and the Top-5 predictions are extracted from the JSON response. 95% confidence intervals are 1,000-replicate bootstrap intervals over the 165 queries. ‘*Δ vs SM-no-alias*’ gives the absolute Top-1 difference against the no-alias baseline. ‘*Δ vs matched-model SM+alias*’ compares each LLM-only row to the *SchemaMapper* hybrid that uses the corresponding alias dictionary. Claude Opus 4.7 has no LLM-only row (see Methods). We used the Top-5 retrievals and the *micro-average* for this evaluation. SM = *SchemaMapper*.

| Method | Top-1 [95% CI] | MRR (micro) | $\Delta$ vs SM-no-alias | $\Delta$ vs matched-model SM+alias |
| --- | --- | --- | --- | --- |
| SM (no alias) | 53.94% | 0.615 | — | — |
| SM + Haiku 4.5 alias | 73.33% | 0.771 | +19.4 pp | — |
| SM + Sonnet 4.5 alias | 66.06% | 0.708 | +12.1 pp | — |
| SM + Opus 4.5 alias | 75.76% | 0.8 | +21.8 pp | — |
| SM + Opus 4.7 alias | 70.30% | 0.73 | +16.4 pp | — |
| SM + Gemini 2.5 Flash alias | 67.88% | 0.726 | +13.9 pp | — |
| SM + Gemini 2.5 Pro alias | 72.73% | 0.759 | +18.8 pp | — |
| Claude Haiku 4.5 | 83.6% [78.2, 89.1] | 0.877 | +29.7 pp | +10.3 pp |
| Claude Sonnet 4.5 | 89.7% [84.8, 94.5] | 0.932 | +35.8 pp | +23.6 pp |
| Claude Opus 4.5 | 86.7% [81.2, 91.5] | 0.916 | +32.8 pp | +10.9 pp |
| Gemini 2.5 Flash | 83.0% [77.0, 88.5] | 0.876 | +29.1 pp | +15.1 pp |
| Gemini 2.5 Pro | 89.1% [84.2, 93.3] | 0.925 | +35.2 pp | +16.4 pp |

**Supplementary Table 11.** Direct memorization probes (E4) across five frontier LLMs. Three complementary probes assess whether each model has memorized the GDC clinical schema. P1 (demographic-node recall): when prompted to enumerate the columns of the GDC demographic node, the percentage of generated lines whose first emitted token matches a valid GDC field name; parentheses report (valid hits / total lines emitted). Haiku 4.5 produced only five generic placeholders rather than schema-grounded column names. P2 (reverse concept → GDC): exact-match rate when the model is given 20 clinical concept descriptions in isolation (no schema context) and asked to return the canonical GDC identifier; parentheses report (correct / 20). P3 (random 3-slot recovery): across ten 12-name slices of the GDC schema, three names per slice are masked at random positions, and the model is asked to recover them; the reported value is blank-level pooled exact-match recall — total correctly recovered names divided by 30 total blanks (10 slices × 3 masks per slice), order-independent within each slice. Higher values across all three probes indicate stronger memorization of GDC schema content; P3 is the most stringent probe, requiring positional recovery rather than recognition.

| Model | P1 (demographic-node recall) | P2 (reverse concept → GDC) | P3 (random 3-slot) |
| --- | --- | --- | --- |
| Claude Haiku 4.5 | 0% (5 unspecific lines) | 65% (13/20) | 0% |
| Claude Sonnet 4.5 | 88% (15/17) | 100% (20/20) | 0% |
| Claude Opus 4.5 | 95% (18/19) | 100% (20/20) | 0% |
| Gemini 2.5 Flash | 83% (5/6) | 65% (13/20) | 0% |
| Gemini 2.5 Pro | 92% (12/13) | 80% (16/20) | 7% |

**Supplementary Table 12.** Matched-control perturbation battery on the GDC benchmark. All configurations are evaluated on the 165-query benchmark (the same as that used in the *SchemaMapper* benchmark) under three perturbations: the original benchmark (E1), source-side variants (E2) reported separately (i.e., synonym-swap and abbreviation-flip), and target-side rename (E3). For ‘*SchemaMapper* (SM)+alias’ under E3, the alias dictionary is translated through the rename map, so aliases continue to point at valid targets. LLM-only baseline values are the names-only configuration. We used the Top-5 retrievals and the micro-average for this evaluation.

| Configuration | E1<br>baseline | E2<br>synonym-swap ($\Delta$) | E2<br>abbrev-flip ($\Delta$) | E3<br>target-rename ($\Delta$) |
| --- | --- | --- | --- | --- |
| SM-no-alias | 53.90% | 33.3% (−20.6) | 35.8% (−18.2) | 40.6% (−13.3) |
| SM + Haiku 4.5 alias | 73.30% | 43.6% (−29.7) | 60.0% (−13.3) | 71.5% (−1.8) |
| SM + Sonnet 4.5 alias | 66.10% | 36.4% (−29.7) | 55.2% (−10.9) | 68.5% (+2.4) |
| SM + Opus 4.5 alias | 75.80% | 50.9% (−24.9) | 59.4% (−16.4) | 78.8% (+3.0) |
| SM + Opus 4.7 alias | 70.30% | 45.5% (−24.8) | 61.2% (−9.1) | 68.5% (−1.8) |
| SM + Gemini 2.5 Flash alias | 67.90% | 42.4% (−25.5) | 52.7% (−15.2) | 70.3% (+2.4) |
| SM + Gemini 2.5 Pro alias | 72.70% | 50.3% (−22.4) | 56.4% (−16.4) | 75.2% (+2.4) |
| LLM-only Haiku 4.5 | 83.60% | 75.2% (−8.5) | 83.2% (−0.4) | 64.2% (−19.4) |
| LLM-only Sonnet 4.5 | 89.70% | 78.2% (−11.5) | 83.2% (−6.5) | 66.1% (−23.6) |
| LLM-only Opus 4.5 | 86.70% | 77.6% (−9.1) | 85.1% (−1.6) | 74.5% (−12.1) |
| LLM-only Gemini 2.5 Flash | 83.00% | 75.2% (−7.9) | 80.7% (−2.3) | 64.8% (−18.2) |
| LLM-only Gemini 2.5 Pro | 89.10% | 80.0% (−9.1) | 85.7% (−3.4) | 71.5% (−17.6) |

**Supplementary Table 13.** LLM-only baselines fall short of *OntologyMapper* on the UKBB–EFO benchmark. The ontology-mapping performance of 888 UKBB-EFO benchmark queries was evaluated across three methods: *OntologyMapper*, LLM-only *zero-shot*, and LLM-only *open_book_full*. The LLM-only approach prompts Claude Haiku 4.5 (temperature = 0) to return the Top-5 EFO labels. For *zero-shot*, no candidate list was provided, and for *open_book_full*, the full EFO label corpus is provided in context. All three methods were evaluated against the same 17,638-label EFO corpus; the 33,230-label EFO corpus used in the *text2term* benchmarking exceeds Haiku 4.5’s input limit and was therefore not used here. ‘Halluc. (Top-1)’ is the fraction of queries whose Top-1 prediction is absent from the corpus. We used the Top-5 retrievals and the micro-average for this evaluation.

| Method | Top-1 (%) | Top-3 (%) | Top-5 (%) | MRR (micro) | Halluc. (Top-1, %) |
| --- | --- | --- | --- | --- | --- |
| *OntologyMapper* | 82.98 | 93.95 | 96.75 | 0.887 | — |
| LLM-only, zero_shot | 54.88 | 61.84 | 63.08 | 0.584 | 34.46 |
| LLM-only, open_book_full | 63.75 | 71.04 | 73.06 | 0.676 | 5.95 |

**Supplementary Table 14.**
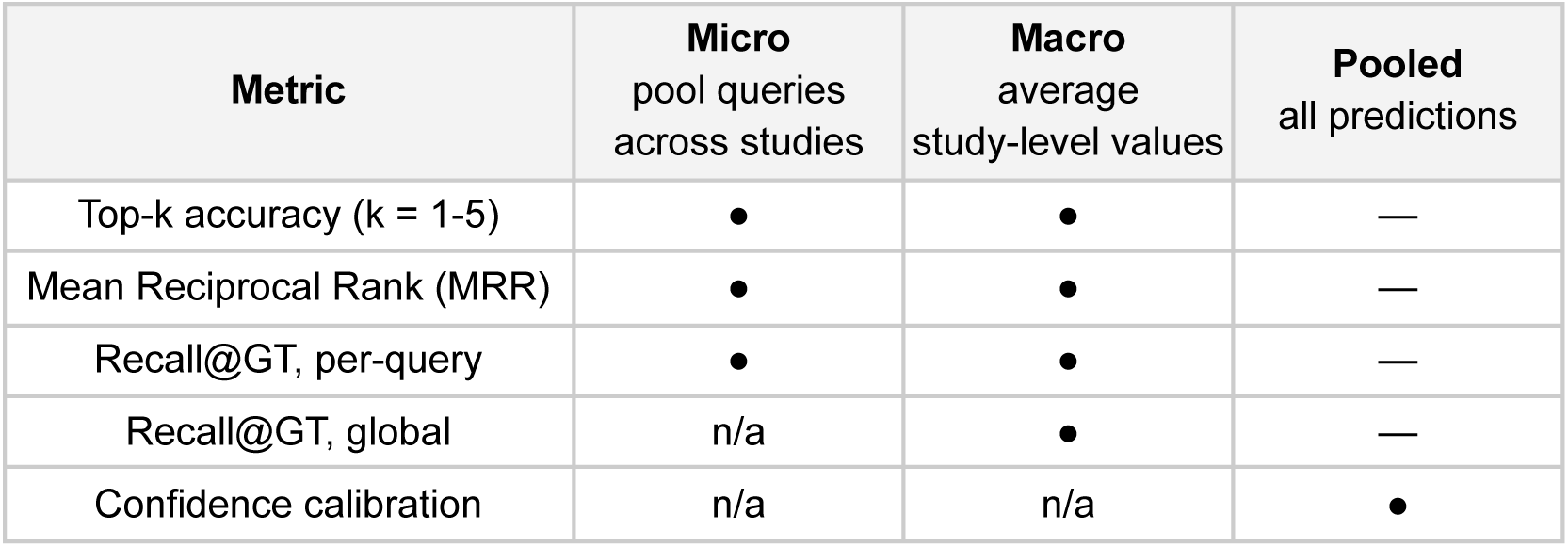
Evaluation metrics and applicable aggregation schemes. Filled circles (●) indicate aggregation schemes reported in this study. Em-dashes (—) indicate schemes that are not applicable because pooled aggregation coincides with micro-averaging for per-query metrics and so is not reported separately. “n/a” indicates schemes that are undefined or degenerate for the metric: micro-averaging is not defined for global Recall@GT, which produces a single study-level value by construction; both micro and macro are degenerate for confidence calibration metrics on small studies. Cross-method comparisons in Table 1 use macro averaging throughout to match the convention used by Liu et al. in the source Magneto evaluation.

| **Metric** | **Micro**<br>pool queries<br>across studies | **Macro**<br>average<br>study-level values | **Pooled**<br>all predictions |
| --- | --- | --- | --- |
| Top-k accuracy (k = 1-5) | ● | ● | — |
| Mean Reciprocal Rank (MRR) | ● | ● | — |
| Recall@GT, per-query | ● | ● | — |
| Recall@GT, global | n/a | ● | — |
| Confidence calibration | n/a | n/a | ● |

**Supplementary Table 15.** Practical guide to LLM model selection for *SchemaMapper* alias generation. Decision-support summary for the seven alias dictionaries evaluated on the GDC schema-matching benchmark, ranked by MRR within performance tiers. For each model, the table reports vendor, hosting mode (API vs. local), MRR with 95% paired-bootstrap CI, Top-1 accuracy, one-time dictionary-generation cost (USD; $0 for locally hosted models), wall-clock generation time, and a recommended use case. Tier-A comprises the six dictionaries statistically equivalent to one another on Top-1 accuracy (all-pairs McNemar, Holm-adjusted over 28 comparisons); within this tier, point estimates favor Opus 4.5, but Haiku 4.5 is statistically indistinguishable from Opus 4.5 on every metric at roughly 20× lower cost and ∼2.8x faster generation time, making it the recommended default. Gemini 2.5 Flash is the lowest-cost option, and Gemini 2.5 Pro is the vendor-diversified choice; Sonnet 4.5 and Opus 4.7 are Pareto-dominated and not recommended for new runs. Tier B (Gemma 3 27B) is a local option with no measurable benefit over no-alias on this benchmark. Tier C is the no-alias baseline. Costs reflect public list prices as of late 2025/early 2026 and exclude batch-API or prompt-caching discounts.

| Method | Vendor | Hosting | MRR | Top-1 (%) | One-time cost | Gen. time | When to choose |
| --- | --- | --- | --- | --- | --- | --- | --- |
| **Tier A (statistically equivalent on MRR / Top-1)** |  |  |  |  |  |  |  |
| Opus 4.5 | Anthropic | API | 0.800 [0.743, 0.855] | 75.76 | \$21.51 | 0.8 h | Max accuracy; cost is no constraint. |
| Haiku 4.5 | Anthropic | API | 0.771 [0.710, 0.828] | 73.33 | \$1.04 | 0.3 h | **Default recommendation — statistically equivalent to Opus 4.5 at ~20× lower cost and ~2.8x faster.** |
| Gemini 2.5 Pro | Google | API | 0.759 [0.695, 0.819] | 72.73 | \$2.00 | 0.7 h | Vendor-diverse alternative to Anthropic at low cost. |
| Opus 4.7 | Anthropic | API | 0.730 [0.664, 0.794] | 70.30 | \$28.13 | 2.5 h | Dominated by Opus 4.5 (same vendor, more expensive, worse MRR). Avoid for new runs. |
| Gemini 2.5 Flash | Google | API | 0.726 [0.662, 0.789] | 67.88 | \$0.49 | 0.2 h | Cheapest API option; equivalent to Tier A on MRR. |
| Sonnet 4.5 | Anthropic | API | 0.708 [0.643, 0.772] | 66.06 | \$4.45 | 0.8 h | Dominated by Haiku 4.5 (same vendor, more expensive, similar MRR). Avoid for new runs. |
| **Tier B (below Tier A)** |  |  |  |  |  |  |  |
| Gemma 3 27B | Google | local | 0.616 [0.546, 0.685] | 56.36 | \$0 (local) | 4.7 h | Air-gapped/on-prem only. No measurable benefit over no-alias on this benchmark. |
| **Tier C (baseline)** |  |  |  |  |  |  |  |
| No alias | NA | NA | 0.615 [0.548, 0.682] | 53.94 | \$0 (none) | — | Baseline only. Tier-A alias can add ~12–19 pp Top-1 at <\$5. |

